# Anatomical connectivity along the anterior-posterior axis of the human hippocampus: new insights using quantitative fibre-tracking

**DOI:** 10.1101/2021.11.17.469032

**Authors:** Marshall A. Dalton, Arkiev D’Souza, Jinglei Lv, Fernando Calamante

## Abstract

The hippocampus supports multiple cognitive functions including episodic memory. Recent work has highlighted functional differences along the anterior-posterior axis of the human hippocampus but the neuroanatomical underpinnings of these differences remain unclear. We leveraged track-density imaging to systematically examine anatomical connectivity between the cortical mantle and the anterior-posterior axis of the *in-vivo* human hippocampus. We first identified the most highly connected cortical areas and detailed the degree to which they preferentially connect along the anterior-posterior axis of the hippocampus. Then, using a tractography pipeline specifically tailored to measure the location and density of streamline endpoints within the hippocampus, we characterised where, within the hippocampus, these cortical areas preferentially connect. Our results were striking in showing that different parts of the hippocampus preferentially connect with distinct cortical areas. Furthermore, we provide evidence that both gradients and circumscribed areas of dense extrinsic anatomical connectivity exist within the human hippocampus. These findings inform conceptual debates in the field by unveiling how specific regions along the anterior-posterior axis of the hippocampus are associated with different cortical inputs/outputs. Overall, our results represent a major advance in our ability to map the anatomical connectivity of the human hippocampus *in-vivo and* inform our understanding of the neural architecture of hippocampal dependent memory systems in the human brain. This detailed characterization of how specific portions of the hippocampus anatomically connect with cortical brain regions may promote a better understanding of its role in cognition and we emphasize the importance of considering the hippocampus as a heterogeneous structure.

## Introduction

There is long-standing agreement that the hippocampus is essential for supporting episodic long-term memory (1) and for facilitating spatial navigation (2). The hippocampus has more recently been linked with other roles including imagination of fictitious and future experiences (3, 4), visuospatial mental imagery (5), visual perception (6, 7) and decision making (8). It is a complex structure containing multiple subregions (referred to as subfields) including the dentate gyrus (DG), cornu ammonis (CA) 4-1, subiculum, presubiculum and parasubiculum. Accumulating evidence suggests that different hippocampal subfields are preferentially recruited during different cognitive functions (5, 9). Functional differences are also present along the anterior-posterior axis of the hippocampus (10–15) and its subfields (16, 17). Despite recent advances in understanding functional differentiation within the hippocampus, much less is known about the neuroanatomical underpinnings of these functional differences in the human brain. A more detailed understanding of anatomical connectivity along the anterior-posterior axis of the human hippocampus is needed to better understand and interpret these functional differences.

Much of our knowledge regarding the anatomical connectivity of the human hippocampus is inferred from the results of tract-tracing studies in rodent and non-human primate brains. While this information has been fundamental to inform theoretical models of hippocampal-dependent memory function, recent investigations have highlighted potential differences between connectivity of the human and non-human primate hippocampus (18). This suggests that, in addition to evolutionarily conserved patterns of hippocampal connectivity, human and non-human primates may also have unique patterns of connectivity. More detailed characterisations of human hippocampal connectivity are, therefore, essential to advance our understanding of the neural architecture that underpins hippocampal dependent memory and cognition in the human brain.

Tract-tracing studies in rodents and non-human primates have revealed that the hippocampus is highly connected with multiple cortical areas. It is well established that the entorhinal cortex (EC) is the primary interface between the hippocampus and multiple brain regions (19). Other medial temporal lobe (MTL) structures including the perirhinal (PeEc) and posterior parahippocampal (PHC) cortices also have direct anatomical connections with the hippocampus (20–23) albeit to a lesser degree. However, direct cortico-hippocampal pathways are not confined to MTL cortices. Anterograde and retrograde labelling studies in non-human primates have revealed direct and reciprocal connections between the hippocampus and multiple cortical areas in temporal (21, 24), parietal (25, 26) and frontal (27, 28) lobes. These detailed investigations show that specific cortical areas preferentially connect with circumscribed portions along the anterior-posterior axis of hippocampal subfields. For example, retrograde labelling studies in the macaque reveal that the retrosplenial cortex (RSC) in the parietal lobe has direct connectivity with posterior portions of the presubiculum while area TE in the inferior temporal lobe displays preferential connectivity with posterior portions of the CA1/subiculum transition area (22). While patterns of cortico-hippocampal connectivity such as these have been observed in the non-human primate brain, we know less about these patterns in the human brain.

Detailed examination of structural connectivity (SC) of the human hippocampus has been difficult to pursue mainly due to the technical difficulties inherent to probing hippocampal connectivity *in-vivo* using MRI. Limitations in both image resolution and fibre tracking methods have precluded our ability to probe cortico-hippocampal pathways in a sufficient level of detail. Some researchers have partially circumvented these constraints by investigating blocks of ex-vivo MTL tissue using high-field MRI scanners (29–31) or novel methods such as polarised light microscopy (18). While these studies have provided important insights relating to MTL-hippocampal pathways, we have less knowledge regarding how the human hippocampus connects with more distant cortical areas. Detailed characterisations of anatomical connectivity between the hippocampal long-axis and broader cortical networks are needed to better understand functional heterogeneity within the hippocampus (32).

Of the extant *in-vivo* MRI studies that have examined SC of the human hippocampus, several have focussed on connectivity between the hippocampus and specific cortical or subcortical areas (33–37) with a primary focus on disease states. To our knowledge, only one study has attempted to characterise the broader hippocampal ‘connectome’ in the healthy human brain. Maller and colleagues (38) used diffusion MRI (dMRI) data with multiple diffusion strengths and high angular resolution combined with track density imaging (TDI) to characterise SC between the whole hippocampus and cortical and subcortical brain regions. A quantitative analysis of streamline numbers (as a surrogate measure of connectivity) seeded from the whole hippocampus showed that the most highly connected brain regions were the temporal lobe followed by subcortical, occipital, frontal and parietal regions. The primary focus of their study, however, was a description of six dominant white matter pathways that accounted for most cortical and subcortical streamlines connecting with the whole hippocampus. Together, these studies have provided an important glimpse into the complexity of human hippocampal SC but important gaps in our knowledge remain. Specifically, there has been no systematic examination of SC between the cortical mantle and the anterior-posterior axis of the human hippocampus and we do not know where, within the hippocampus, specific cortical areas preferentially connect.

In the current study, we aimed to systematically examine patterns of SC between cortical brain areas and the anterior-posterior axis of the human hippocampus. We combined high-quality dMRI data from the Human Connectome Project (HCP), cutting-edge quantitative fibre-tracking methods, and a processing pipeline specifically tailored to study hippocampal connectivity with three primary aims; (i) to quantitatively characterise SC between the cortical mantle (focused on non-MTL areas) and the whole hippocampus; (ii) to quantitatively characterise how SC varies between cortical areas and the head, body and tail of the hippocampus and (iii) to use TDI combined with ‘endpoint density mapping’ to quantitatively assess, visualise and map the spatial distribution of streamline endpoints within the hippocampus associated with each cortical area.

Our results represent a major advance in; (i) our ability to map the anatomical connectivity of the human hippocampus *in-vivo* and; (ii) our understanding of the neural architecture that underpins hippocampal dependent memory systems in the human brain. We provide fundamental insights into how specific cortical areas preferentially connect along the anterior-posterior axis of the hippocampus and identify where streamlines associated with a given cortical area preferentially connect within the hippocampus. These detailed anatomical insights will help fine-tune network connectivity models and will have an impact on current theoretical models of human hippocampal memory function^1^.

## Results

We first characterised SC between the whole hippocampus and all cortical areas of the HCP Multi-Modal Parcellation (HCPMMP) (39). The primary focus of this study was SC between the hippocampus and non-MTL cortical areas. We therefore focus on cortical areas outside of the MTL (information relating to MTL cortices are presented in Supplementary Materials Note S1, Supplementary Figures S1 and S2 and Supplementary Tables S1 and S2).

### Which specific cortical areas most strongly connect with the whole hippocampus?

For brevity, we present results relating to the twenty cortical areas with the highest degree of SC with the whole hippocampus. The hippocampus displayed the highest degree of SC with discrete cortical areas in temporopolar (areas TGv, TGd; abbreviations for all cortical areas are defined in Supplementary Table S3), inferolateral temporal (areas TF, TE2a, TE2p), medial parietal (areas RSC, ProS, POS1, POS2, DVT) dorsal and ventral stream visual (areas V3A, V6, FFC, VVC, VMV1, VMV2) and early visual (occipital) cortices (areas V1, V2, V3, V4). Results are summarised in Figure 1a and Table 1 lists each cortical area and their associated strength of connectivity with the whole hippocampus. A full list of all cortical areas of the HCPMMP and their associated strengths of connectivity are provided in Supplementary Table S1.

**Figure 1.**
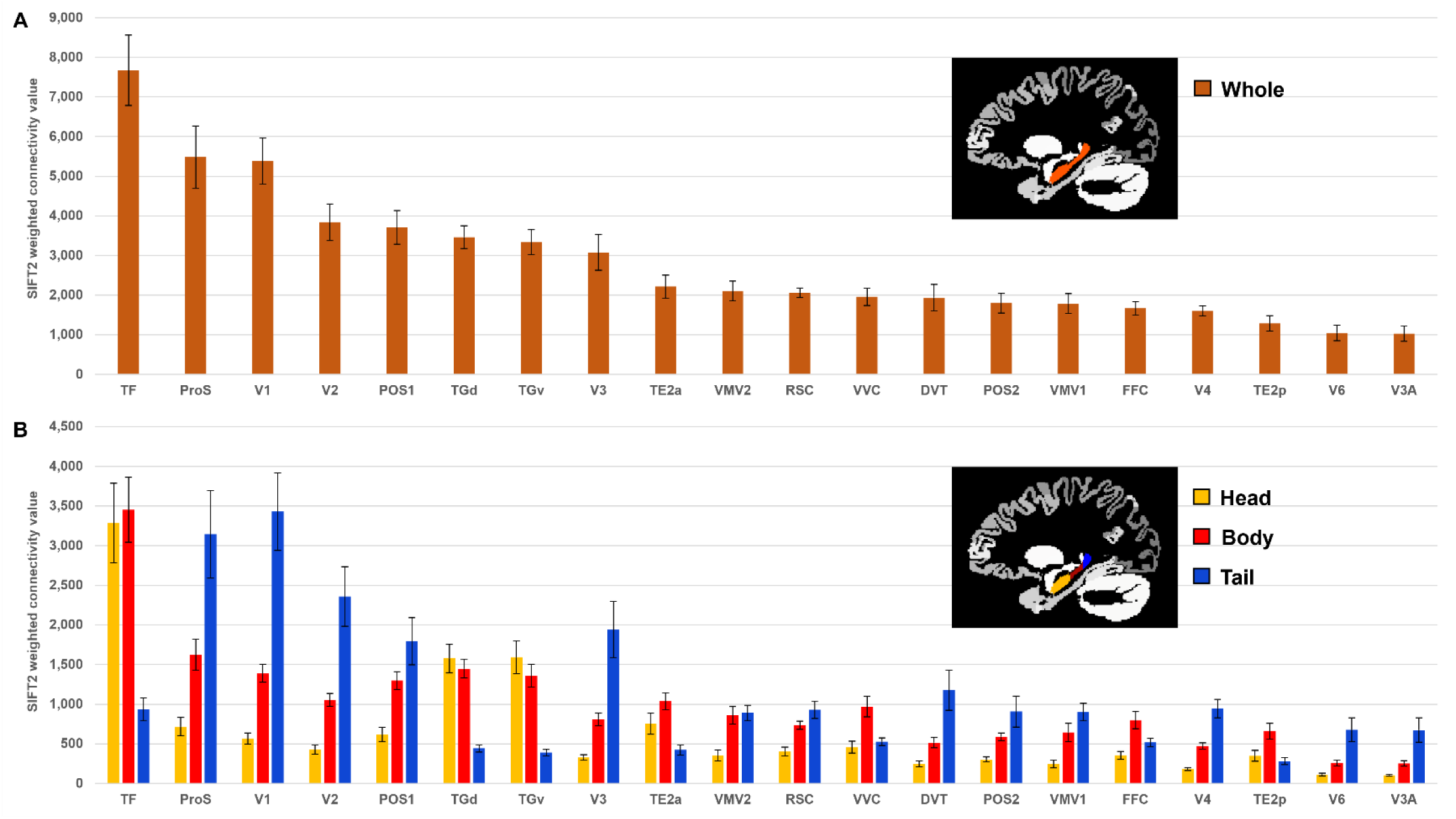
Twenty cortical brain areas with the highest degree of anatomical connectivity with the hippocampus. a. Histogram plotting the mean structural connectivity (given by the sum of SIFT2 weighted values) associated with the twenty cortical areas most strongly connected with the whole hippocampus (excluding medial temporal lobe (MTL) areas; see Supplementary Figure S1 for MTL values). Error bars represent the standard error of the mean. b. Histogram plotting the corresponding mean SIFT2 weighted values associated with anterior (yellow), body (red) and tail (blue) portions of the hippocampus for the twenty most strongly connected cortical areas presented in a. Errors bars represent the standard error of the mean.

**Table 1.** Twenty cortical brain areas (excluding medial temporal lobe) with the highest degree of anatomical connectivity with the whole hippocampus. Column 1 displays cortical areas as defined by the Human Connectome Project Multi-Modal Parcellation (HCPMMP) scheme and ordered by strength of connectivity with the whole hippocampus (abbreviations for all cortical areas are defined in Supplementary Table S3). Column 2 indicates the broader brain region within which each cortical area is located. Column 3 displays the mean SIFT2 weighted value (connectivity strength) associated with each brain area. Column 4 displays the standard error of the mean. Column 5 displays the percent of all cortical connections accounted for by each area. Values in brackets indicate the percent of cortical connections accounted for by each area when excluding medial temporal lobe (MTL) areas.

| Cortical area | Location of area | Whole hippocampus |  |  |
| --- | --- | --- | --- | --- |
|  |  | Mean SIFT2 weighted value (connectivity strength; n=10) | SE of mean | Percent of all cortical connections accounted for by area. (Percent of cortical connections excluding MTL areas) |
| TF | Lateral Temporal Cortex | 7673 | 886 | 5.10 (10.61) |
| ProS | Medial Parietal Cortex (including Posterior Cingulate) | 5483 | 784 | 3.64 (7.58) |
| V1 | Early Visual Cortex (Occipital) | 5385 | 579 | 3.58 (7.45) |
| V2 | Early Visual Cortex (Occipital) | 3840 | 462 | 2.55 (5.31) |
| POS1 | Medial Parietal Cortex (including Posterior Cingulate) | 3712 | 424 | 2.47 (5.13) |
| TGd | Temporal Pole | 3465 | 288 | 2.30 (4.79) |
| TGv | Temporal Pole | 3337 | 313 | 2.22 (4.61) |
| V3 | Early Visual Cortex (Occipital) | 3079 | 450 | 2.05 (4.26) |
| TE2a | Lateral Temporal Cortex | 2214 | 288 | 1.47 (3.06) |
| VMV2 | Ventral Stream Visual Cortex | 2105 | 247 | 1.40 (2.91) |
| RSC | Medial Parietal Cortex (including Posterior Cingulate) | 2063 | 121 | 1.37 (2.85) |
| VVC | Ventral Stream Visual Cortex | 1956 | 220 | 1.30 (2.71) |
| DVT | Medial Parietal Cortex (including Posterior Cingulate) | 1939 | 335 | 1.29 (2.68) |
| POS2 | Medial Parietal Cortex (including Posterior Cingulate) | 1802 | 248 | 1.20 (2.49) |
| VMV1 | Ventral Stream Visual Cortex | 1788 | 248 | 1.19 (2.47) |
| FFC | Ventral Stream Visual Cortex | 1670 | 172 | 1.11 (2.31) |
| V4 | Early Visual Cortex (Occipital) | 1601 | 127 | 1.06 (2.21) |
| TE2p | Lateral Temporal Cortex | 1288 | 198 | 0.86 (1.78) |
| V6 | Dorsal Stream Visual Cortex | 1050 | 195 | 0.70 (1.45) |
| V3A | Dorsal Stream Visual Cortex | 1029 | 197 | 0.68 (1.42) |

### Do cortical areas display preferential connectivity along the anterior-posterior axis of the hippocampus?

Next, we conducted a more detailed characterisation of SC between each cortical area and the head, body and tail portions of the hippocampus. For brevity, we present results relating to the twenty most highly connected cortical areas described above. Results are summarised in Figure 1b and Table 2 which lists each cortical area, their associated strength of connectivity with the head, body and tail portions of the hippocampus and the results of statistical analyses (Bonferroni corrected paired-samples t-tests; see Methods). A full list of all cortical areas and their associated strengths of connectivity with the head, body and tail portions of the hippocampus are provided in Supplementary Table S2.

**Table 2.** Connectivity between the twenty most highly connected cortical areas and the head, body and tail of the hippocampus. Column 1 displays cortical areas as defined by the Human Connectome Project Multi-Modal Parcellation (HCPMMP) scheme and ordered by strength of connectivity with the whole hippocampus (abbreviations for all cortical areas are defined in Supplementary Table S3). Column 2 designates the portion of hippocampus (head, body, tail). Column 3 displays the mean SIFT2 weighted value (connectivity strength) between each cortical area and the head, body and tail of the hippocampus. Column 4 displays the standard error of the mean. Column 5 displays the contrast for each paired samples t-test. Column 6 displays the t-statistic associated with each pair. Column 7 displays the p-value associated with each pair. Column 8 indicates the significance level associated with each pair following Bonferroni correction. *** = <0.001, ** = <0.01, * = <0.05, n.s. = not statistically significant.

| Cortical area | Portion of hippocampus | Mean SIFT2 weighted value (connectivity strength; n=10) | SE of Mean | Paired samples t-test |  |  |  |
| --- | --- | --- | --- | --- | --- | --- | --- |
|  |  |  |  | Pair | t-statistic t(9) | p-value | Significance |
| TF | Head | 3287 | 500 | Head - body | -0.34 | 0.741 | n.s. |
|  | Body | 3452 | 410 | Body - tail | 9.077 | <0.001 | *** |
|  | Tail | 934 | 144 | Head - tail | 5.049 | 0.001 | ** |
| ProS | Head | 716 | 116 | Head - body | -5.75 | <0.001 | *** |
|  | Body | 1625 | 194 | Body - tail | -3.618 | 0.006 | ** |
|  | Tail | 3143 | 553 | Head - tail | -4.799 | 0.001 | ** |
| V1 | Head | 567 | 70 | Head - body | -9.508 | <0.001 | *** |
|  | Body | 1389 | 112 | Body - tail | -4.629 | 0.001 | ** |
|  | Tail | 3429 | 489 | Head - tail | -5.919 | <0.001 | *** |
| V2 | Head | 428 | 62 | Head - body | -11.910 | <0.001 | *** |
|  | Body | 1054 | 82 | Body - tail | -3.903 | 0.004 | ** |
|  | Tail | 2358 | 376 | Head - tail | -5.396 | <0.001 | *** |
| POS1 | Head | 620 | 89 | Head - body | -6.255 | <0.001 | *** |
|  | Body | 1299 | 112 | Body - tail | -2.196 | 0.056 | n.s. |
|  | Tail | 1793 | 297 | Head - tail | -4.043 | 0.003 | ** |
| TGd | Head | 1576 | 182 | Head - body | 0.79 | 0.450 | n.s. |
|  | Body | 1446 | 117 | Body - tail | 11.378 | <0.001 | *** |
|  | Tail | 443 | 47 | Head - tail | 6.641 | <0.001 | *** |
| TGv | Head | 1589 | 207 | Head - body | 1.121 | 0.291 | n.s. |
|  | Body | 1358 | 144 | Body - tail | 8.211 | <0.001 | *** |
|  | Tail | 389 | 41 | Head - tail | 5.973 | <0.001 | *** |
| V3 | Head | 327 | 37 | Head - body | -7.204 | <0.001 | *** |
|  | Body | 810 | 83 | Body - tail | -3.921 | 0.004 | ** |
|  | Tail | 1942 | 357 | Head - tail | -4.718 | 0.001 | ** |
| TE2a | Head | 754 | 132 | Head - body | -3.759 | 0.004 | ** |
|  | Body | 1038 | 107 | Body - tail | 11.479 | <0.001 | *** |
|  | Tail | 422 | 65 | Head - tail | 3.787 | 0.004 | ** |
| VMV2 | Head | 354 | 68 | Head - body | -7.919 | <0.001 | *** |
|  | Body | 860 | 111 | Body - tail | -0.375 | 0.716 | n.s. |
|  | Tail | 891 | 97 | Head - tail | -6.248 | <0.001 | *** |
| RSC | Head | 401 | 55 | Head - body | -8.647 | <0.001 | *** |
|  | Body | 733 | 52 | Body - tail | -1.55 | 0.155 | n.s. |
|  | Tail | 928 | 107 | Head - tail | -3.727 | 0.005 | ** |
| VVC | Head | 460 | 76 | Head - body | -5.554 | <0.001 | *** |
|  | Body | 969 | 129 | Body - tail | 4.058 | 0.003 | ** |
|  | Tail | 528 | 48 | Head - tail | -0.885 | 0.399 | n.s. |
| DVT | Head | 247 | 36 | Head - body | -5.436 | <0.001 | *** |
|  | Body | 515 | 65 | Body - tail | -3.374 | 0.008 | ** |
|  | Tail | 1178 | 253 | Head - tail | -3.954 | 0.003 | ** |
| POS2 | Head | 303 | 32 | Head - body | -6.682 | <0.001 | *** |
|  | Body | 590 | 47 | Body - tail | -2.02 | 0.074 | n.s. |
|  | Tail | 909 | 195 | Head - tail | -3.211 | 0.011 | * |
| VMV1 | Head | 245 | 47 | Head - body | -5.392 | <0.001 | *** |
|  | Body | 644 | 115 | Body - tail | -2.939 | 0.017 | n.s. (following<br>Bonferroni correction) |
|  | Tail | 899 | 110 | Head - tail | -7.372 | <0.001 | *** |
| FFC | Head | 355 | 49 | Head - body | -5.364 | <0.001 | *** |
|  | Body | 798 | 109 | Body - tail | 2.805 | 0.021 | n.s. (following Bonferroni correction) |
|  | Tail | 517 | 55 | Head - tail | -2.24 | 0.052 | n.s. |
| V4 | Head | 183 | 20 | Head - body | -10.472 | <0.001 | *** |
|  | Body | 472 | 40 | Body - tail | -4.002 | 0.003 | ** |
|  | Tail | 946 | 116 | Head - tail | -6.136 | <0.001 | *** |
| TE2p | Head | 348 | 69 | Head - body | -5.453 | <0.001 | *** |
|  | Body | 658 | 101 | Body - tail | 5.168 | 0.001 | ** |
|  | Tail | 282 | 45 | Head - tail | 1.163 | 0.275 | n.s. |
| V6 | Head | 113 | 16 | Head - body | -5.428 | <0.001 | *** |
|  | Body | 259 | 37 | Body - tail | -3.652 | 0.005 | ** |
|  | Tail | 678 | 149 | Head - tail | -4.047 | 0.003 | ** |
| V3A | Head | 103 | 11 | Head - body | -5.641 | <0.001 | *** |
|  | Body | 252 | 35 | Body - tail | -3.46 | 0.007 | ** |
|  | Tail | 674 | 155 | Head - tail | -3.895 | 0.004 | ** |

Each of the twenty most highly connected cortical areas displayed preferential connectivity with specific regions along the anterior-posterior axis of the hippocampus. These can be categorised into four distinct patterns: 1. an anterior-to-posterior gradient of increasing connectivity; 2. a posterior connectivity bias; 3. an anterior connectivity bias; and 4. a body connectivity bias.

Eight of the twenty most highly connected cortical areas displayed a gradient of increasing connectivity from the head-to-tail of the hippocampus. These were area ProS, V1, V2, V3, DVT, V4, V6 and V3A (abbreviations are defined in Supplementary Table S3). The results of Bonferroni corrected paired-samples t-tests revealed that these regions each showed a statistically significant difference in connectivity strength between the head and body, the body and tail and the head and tail portions of the hippocampus (see Figure 1b and Table 2). That is, each of these cortical areas displayed the lowest connectivity with the hippocampal head, significantly increased connectivity with the body and the strongest connectivity with the tail.

Five areas had significantly stronger connectivity with the posterior 2/3 of the hippocampus. These were area POS1, VMV1, VMV2, RSC and POS2. The results of Bonferroni corrected paired-samples t-tests revealed that these cortical areas displayed a high degree of connectivity with the body and tail of the hippocampus (no statistically significant difference in connectivity) and significantly lower connectivity with the hippocampal head (see Figure 1b and Table 2).

Three of the twenty most highly connected cortical areas had significantly greater connectivity with the anterior 2/3 of the hippocampus These were area TF, TGd and TGv. The results of Bonferroni corrected paired-samples t-tests revealed that these cortical areas displayed a high degree of connectivity with the head and body of the hippocampus (no statistically significant difference in connectivity) and significantly lower connectivity with the tail of the hippocampus (see Figure 1b and Table 2).

Four areas had greater connectivity with the body of the hippocampus. These areas were TE2a, VVC, FFC and TE2p. The results of Bonferroni corrected paired-samples t-tests revealed that these areas showed statistically significant differences in connectivity strength between the head and body of the hippocampus. TE2a, VVC and TE2p showed statistically significant differences in connectivity strength between the body and tail portions but FFC did not (following Bonferroni correction). TE2a also showed a statistically significant difference between the head and tail portions of the hippocampus but VVC, FFC and TE2p did not (see Figure 1b and Table 2). That is, each of these cortical areas displayed the highest degree of connectivity with the body of the hippocampus.

To summarise, our results detail the degree to which specific cortical areas preferentially connect along the anterior-posterior axis of the hippocampus. While some cortical areas displayed gradients of connectivity strength along the anterior-posterior axis of the hippocampus, others displayed preferential connectivity with specific portions of the hippocampus.

### Do cortical areas display unique distributions of endpoint density within the hippocampus?

Our tractography pipeline was specifically tailored to allow streamlines to enter/leave the hippocampus and quantitatively measure the location and density of streamline endpoints within the hippocampus using TDI. We created endpoint density maps (EDMs) that allowed us to visualise the spatial distribution of hippocampal endpoint density associated with each cortical area (described in Methods). While it is not feasible to present the results for all cortical areas of the HCPMMP, we describe results for the twenty most highly connected cortical areas described above.

The results of group level analyses confirmed that specific cortical areas preferentially connect with different regions within the human hippocampus. For example, areas in the medial parietal cortex (ProS, POS1, RSC, DVT, POS2) displayed high endpoint density primarily in medial portions of the posterior hippocampus (see yellow arrows in Figure 2a for a representative example of endpoint densities associated with RSC; see Supplementary Figure S3 for other areas). In contrast, areas in temporopolar and inferolateral temporal cortex (TF, TGd, TGv, TE2a, TE2p) displayed high endpoint density primarily along the lateral aspect of the anterior 2/3 of the hippocampus and in a circumscribed region of the anterior medial hippocampus (see blue and white arrows respectively in Figure 2b for a representative example of endpoint densities associated with TGv; see Supplementary Figure S4 for other areas). Similar to areas in the medial parietal cortex, areas in the occipital cortices (V1-4, V6, V3a) displayed high endpoint density primarily in the posterior medial hippocampus and, to a lesser degree, in a circumscribed region of the anterior medial hippocampus (see yellow and white arrows respectively in Figure 2c for endpoint densities associated with V1; see Supplementary Figure S5 for other areas).

**Figure 2.**
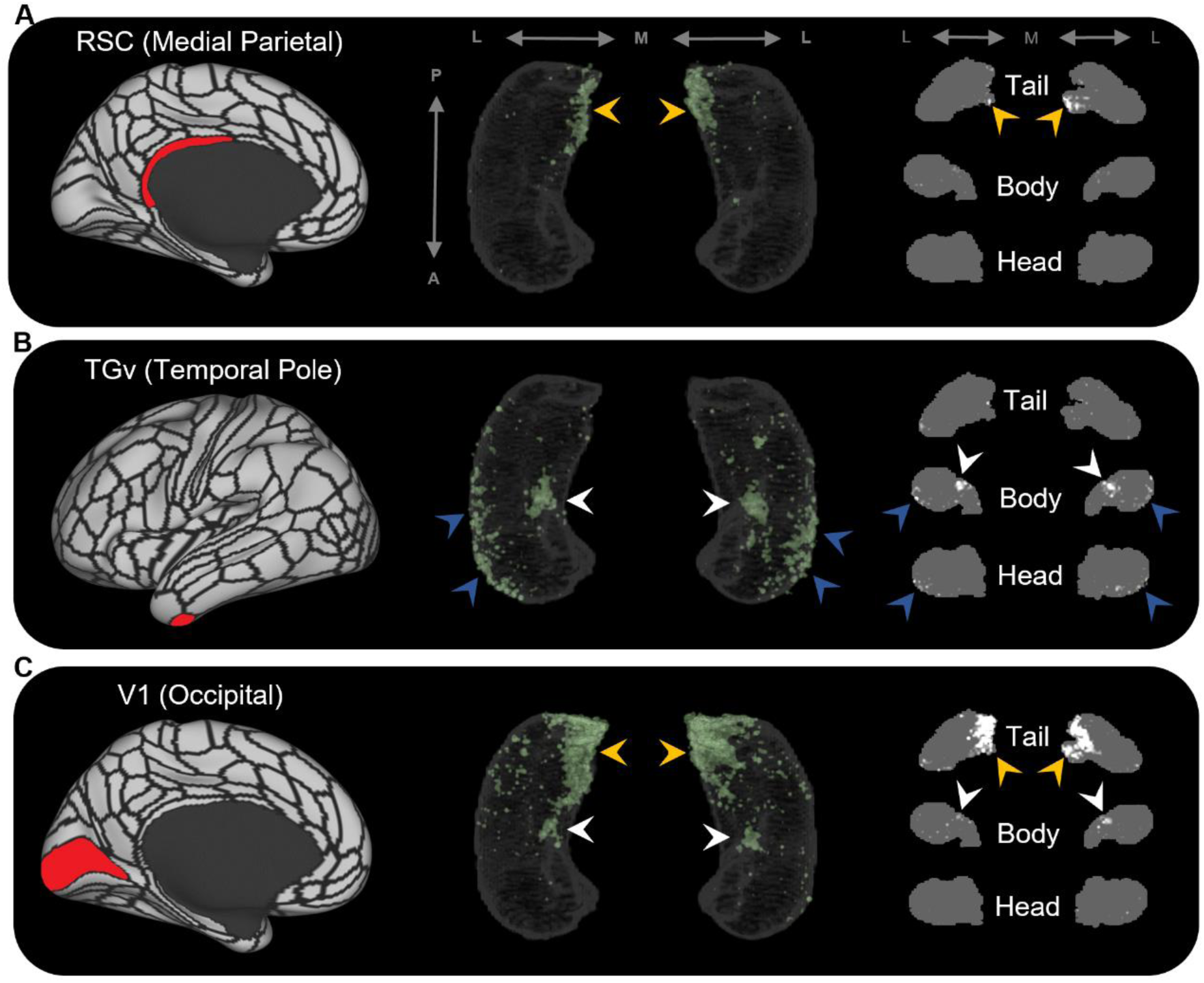
Representative examples of the spatial distribution of endpoint density within the hippocampus for different cortical brain areas. Representative examples of the location of endpoint densities associated with RSC in the medial parietal lobe (a), TGv in the temporal pole (b) and V1 in the occipital lobe (c). In each panel, the location of the relevant brain area is indicated in red on the brain map (left); a 3D rendered representation of the bilateral group level hippocampus mask is presented (middle; transparent grey) overlaid with the endpoint density map associated with each brain area (green); representative slices of the head, body and tail of the hippocampus are displayed in the coronal plane (right; grey) and overlaid with endpoint density maps (white). Note, the spatial distribution of endpoint density within the hippocampus associated with each brain area differs along both the anterior-posterior and medial-lateral axes of the hippocampus. RSC and V1 displayed greatest endpoint density in the posterior medial hippocampus (yellow arrows in a and c). In contrast, TGv displayed greatest endpoint density in the anterior lateral hippocampus and in a circumscribed region in the anterior medial hippocampus (blue and white arrows respectively in b). Area V1 also expressed endpoint density in a circumscribed region in the anterior medial hippocampus (white arrows in c). A=anterior; P=posterior; M=medial; L=lateral.

In parallel with these differences, specific regions within the hippocampus displayed high endpoint density for multiple cortical areas. For example, several medial parietal and occipital cortical areas displayed high endpoint density in the posterior medial hippocampus (yellow arrows in Supplementary Figures S3 and S5). In contrast, several cortical areas in the temporal pole and inferolateral temporal lobe displayed high endpoint density in the anterior lateral hippocampus (blue arrows Supplementary Figure S4). Another cluster of high endpoint density in the anterior medial hippocampus (at the level of the uncal apex) was more broadly associated with specific areas in temporal, medial parietal and occipital cortices (white arrows in Supplementary Figures S3-5).

To further probe hippocampal endpoint density common to these cortical regions, we averaged the EDMs for anatomically related cortical areas. For example, we averaged the EDMs of the five most highly connected cortical areas in the temporal lobe (TF, TGd, TGv, TE2a, TE2p) and observed high endpoint density common to these areas was more clearly localised along the anterior lateral hippocampus and in a circumscribed region of the anterior medial hippocampus (blue and white arrows respectively in Figure 3a). Likewise, when we averaged the most highly connected cortical areas in the medial parietal and occipital cortices respectively, high endpoint density common to each of these areas was localised in the posterior medial hippocampus and, to a lesser degree, in a circumscribed region of the anterior medial hippocampus (Yellow and white arrows respectively in Figure 3b and 3c). When we averaged the EDMs across all of these areas, high endpoint density common to this broader collection was localised to separate circumscribed clusters in the posterior and anterior medial hippocampus (yellow and white arrows respectively in Figure 3d) and in punctate clusters along the anterior-posterior extent of the lateral hippocampus (blue arrows in Figure 3d). Our results suggest that these specific regions within the hippocampus are highly connected with multiple cortical areas in medial parietal, temporal and occipital lobes.

**Figure 3.**
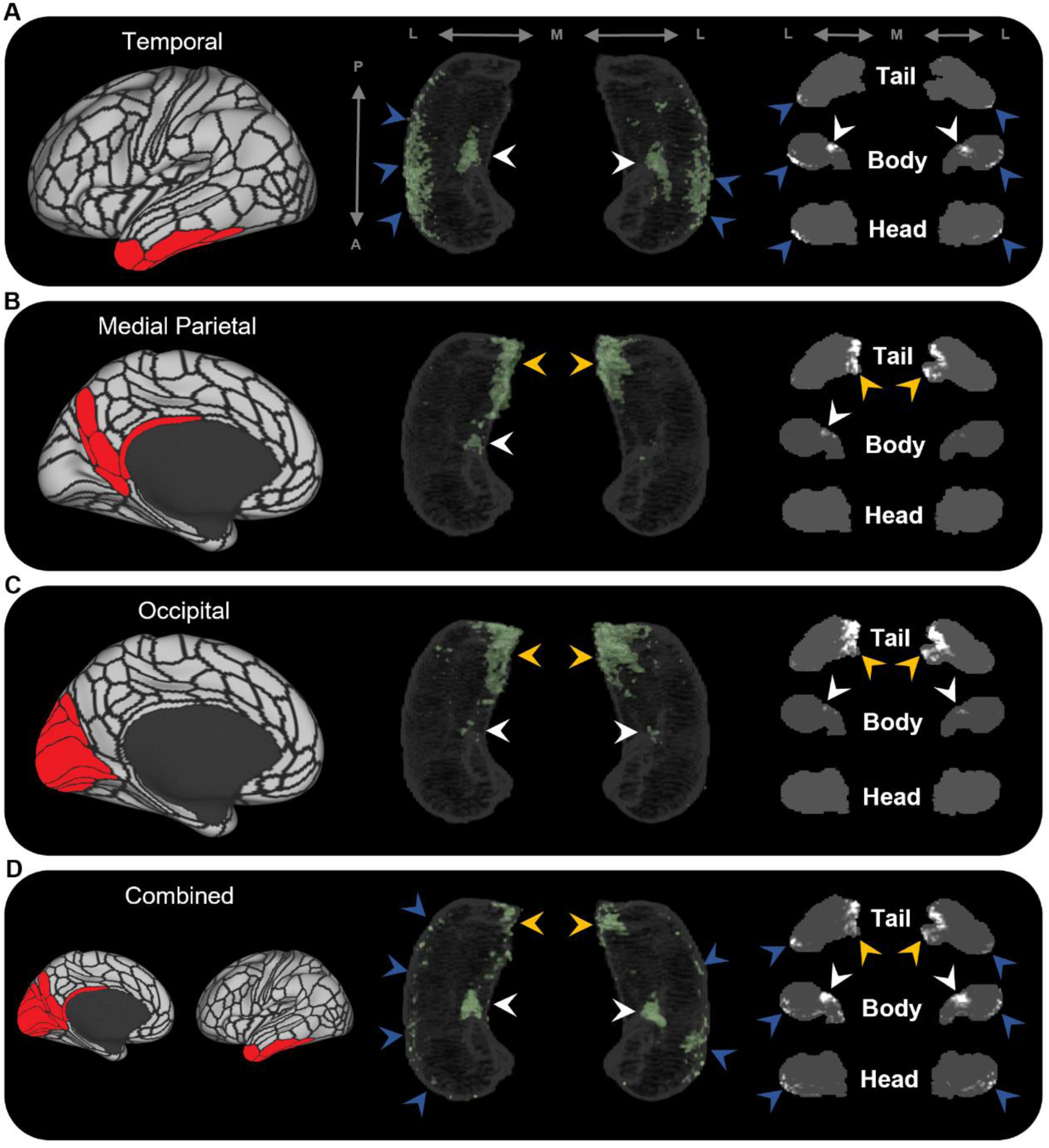
Averaged endpoint density maps for anatomically related brain areas. We averaged the endpoint density maps for the mostly highly connected brain areas in temporal (a; TF, TGd, TGv, TE2a, TE2p), medial parietal (b; ProS, POS1, RSC, DVT, POS2) and occipital (c; V1-4, V6, V3a) cortices and each of these regions combined (d). In each panel, the location of the relevant brain areas are indicated in red on the brain map (left); a 3D rendered representation of the bilateral group level hippocampus mask is presented (middle; transparent grey) overlaid with the endpoint density map associated with each collection of brain areas; representative slices of the head, body and tail of the hippocampus are displayed in the coronal plane (right; grey) and overlaid with endpoint density maps (white). Average endpoint density associated with temporal areas (a) was primarily localised along the anterior lateral hippocampus and a circumscribed region in the anterior medial hippocampus (blue and white arrows respectively). Average endpoint density associated with medial parietal (b) and occipital (c) areas was primarily localised to the posterior medial hippocampus (yellow arrows) and, to a lesser degree, circumscribed regions in the anterior medial hippocampus (white arrows). Average endpoint density associated with these temporal, medial parietal and occipital brain areas combined (d) was localised to circumscribed regions in the posterior and anterior medial hippocampus (yellow and white arrows respectively) and in punctate clusters along the anterior-posterior extent of the lateral hippocampus (blue arrows) suggesting that these specific regions within the hippocampus are highly connected with multiple cortical areas. A=anterior; P=posterior; M=medial; L=lateral.

In addition, our novel method allowed us to isolate and visualise streamlines between specific cortical areas and the hippocampus and map the spatial distribution of hippocampal endpoint density at the single participant level. Representative examples are presented in Figure 4. Figure 4b displays isolated streamlines associated with areas TF and V1 in a single participant. Figure 4d displays the hippocampal EDMs associated with each of these cortical areas in the same participant. When overlaid on the T1-weighted image, EDMs resemble histological staining of post-mortem tissue, albeit *in-vivo* and at a coarser level of detail (see bottom panels of Figure 4d). For example, for area TF, in a coronal slice taken at the uncal apex (red line) endpoint density was primarily localised to specific areas in the medial hippocampus (red panel). For area V1, in a coronal slice taken at the hippocampal tail (turquoise line) endpoint density was also localised to the medial hippocampus (turquoise panel). When compared with equivalent sections of histologically stained tissue, the location of these clusters of endpoint density roughly align with the location of the distal subiculum/proximal presubiculum for both TF and V1 (indicated by black arrows; see also Figure 5a). The location of endpoint density associated with each cortical area was broadly consistent across participants (evidenced by the results of our group level analyses). A detailed examination of individual variability in these patterns, however, was beyond the scope of the current investigation and will be addressed in future studies.

**Figure 4.**
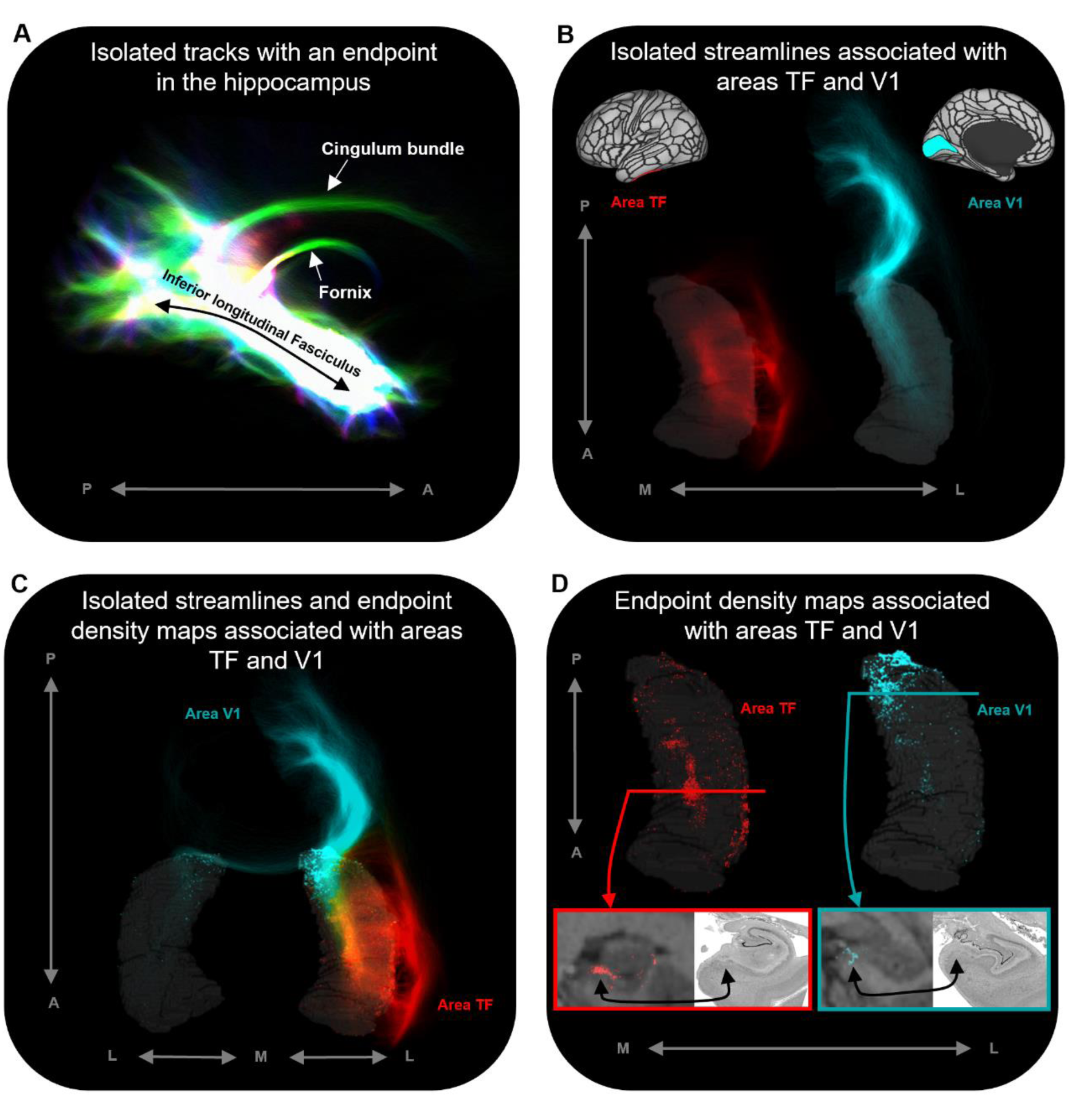
Representative examples of single subject analysis. a. 3D rendering of the hippocampus tractogram for a single participant showing isolated tracks with an endpoint in the hippocampus viewed in the sagittal plane (displayed with transparency; high intensity represents high density of tracks). b. 3D rendered left hippocampus masks (transparent grey) for the same participant overlaid with isolated streamlines associated with the left hemisphere areas TF (red) and V1 (turquoise). The location of areas TF and V1 are indicated on the brain maps (top). c. 3D rendered bilateral hippocampus mask for the same participant (transparent grey) overlaid with isolated streamlines and endpoint density maps associated with the left hemisphere areas TF (red) and V1 (turquoise). Note, while streamlines associated with areas TF and V1 are primarily ipsilateral in nature, streamlines associated with V1 also project to the contralateral hippocampus. d. 3D rendered left hippocampus masks (transparent grey) for the same participant overlaid with endpoint density maps associated with areas TF (red) and V1 (turquoise). For TF and V1, we present a coronal section of the T1-weighted structural image overlaid with the endpoint density maps and a corresponding slice of post-mortem hippocampal tissue (from a different subject) for anatomical comparison (bottom). For both TF (red border; level of the uncal apex) and V1 (turquoise border; level of the hippocampal tail), endpoint density is primarily localised to a circumscribed region in the medial hippocampus aligning with the location of the distal subiculum/proximal presubiculum (black arrows; also see Figure 5a). Post-mortem images are from the BigBrain Project (80). A=anterior; P=posterior; M=medial; L=lateral.

**Figure 5.**
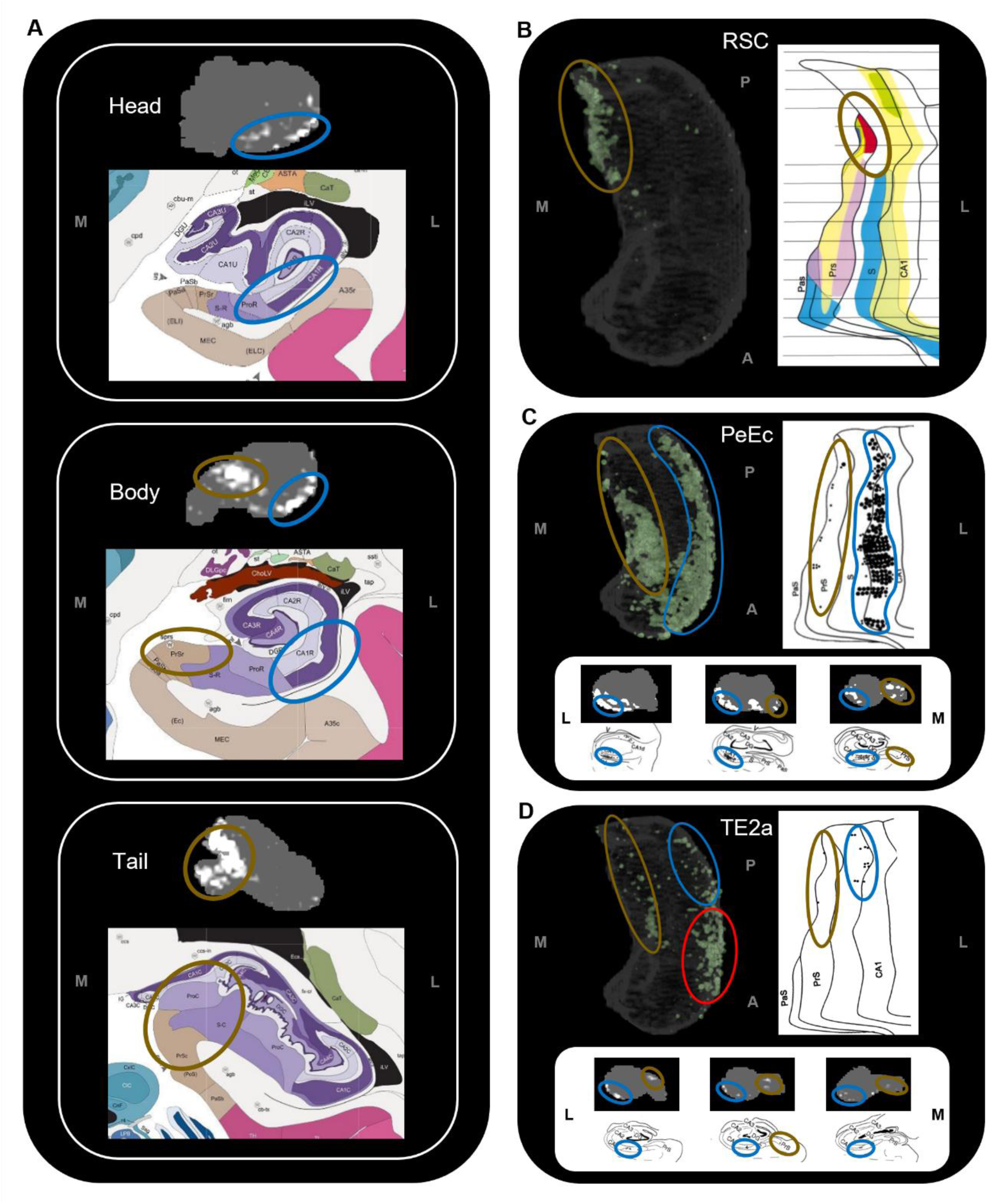
Anatomical location of endpoint densities within the hippocampus and comparison with results of non-human primate studies. a. Representative slices of the head (top), body (middle) and tail (bottom) of the hippocampus displayed in the coronal plane (grey) and overlaid with group level endpoint density maps associated with area TF (head and body; white) and V1 (tail; white). Schematic representations of roughly equivalent slices of the hippocampus showing hippocampal subfields are displayed below each slice (from the Allen Adult Human Brain Atlas website: https://atlas.brain-map.org/). Note, endpoint density in the lateral hippocampus (blue ellipsoids) aligns with the location of the distal CA1/proximal subiculum. Endpoint density in the medial hippocampus (brown ellipsoids) aligns with the location of distal subiculum/proximal presubiculum. b-d. 3D rendered representations of the group level hippocampus mask (left; transparent grey) are overlaid with endpoint density maps (green) associated with RSC (b), PeEc (c) and TE2a (d). Schematic representations of the macaque hippocampus (right; Images reproduced with permission from (22)) show the location of labelled cells following retrograde tracer injection into the RSC (b; red), PeEc (c; black points) and TE2a (d; black points). The bottom panel of c and d displays slices of human hippocampus in the coronal plane (grey) overlaid with endpoint density maps (white) and roughly equivalent slices of the macaque hippocampus display the location of labelled cells (black points; Images reproduced with permission from (22)). Note, endpoint density and labelled cells in the human and macaque respectively are localised to similar regions along the anterior-posterior, lateral (blue ellipsoids) and medial (brown ellipsoids) hippocampus. However, areas of difference also exist (d; red ellipsoid). M=medial; L=lateral; A=anterior; P=posterior.

## Discussion

This study represents a comprehensive *in-vivo* characterisation of SC between cortical brain areas and the human hippocampus. We identified cortical areas with the highest degree of SC with the whole hippocampus, measured the degree to which these cortical areas preferentially connect along the anterior-posterior axis of the hippocampus, and deployed a tailored method to characterise where, within the hippocampus, each cortical area preferentially connects. Our results reveal how specific cortical areas preferentially connect with circumscribed regions along the anterior-posterior and medial-lateral axes of the hippocampus. Our results broadly reflect observations from the non-human primate literature (discussed below) and contribute new neuroanatomical insights to inform debates on human hippocampal function as it relates to its anterior-posterior axis. This work represents an important advance in our understanding of the neural architecture that underpins hippocampal dependent memory systems in the human brain. In addition, our method represents a novel approach to conduct detailed *in-vivo* investigations of the anatomical connectivity of the human hippocampus with implications for basic and clinical neuroscience.

### Preliminary whole hippocampus and anterior-posterior axis analyses

First, we consider the preliminary whole hippocampus analysis. Aligning with our predictions, the hippocampus was most strongly connected with the EC and highly connected with surrounding MTL structures (See Supplementary Materials for further information relating to MTL structures). Beyond the MTL, specific cortical areas in temporopolar, inferotemporal, medial parietal and occipital cortices displayed the highest degree of SC with the hippocampus. These results broadly align with recent DWI investigations that reported SC between the whole hippocampus and these cortical regions in the human brain (38). Our anterior-posterior axis analyses provided a more detailed quantitative characterisation of cortico-hippocampal SC by measuring the degree to which specific cortical areas preferentially connect along the anterior-posterior axis of the hippocampus. In brief, specific areas within temporopolar and inferolateral temporal cortices displayed preferential SC with the head and/or body of the hippocampus. In contrast, medial parietal and occipital cortical areas displayed a posterior hippocampal connectivity bias most strongly with the hippocampal tail. These patterns of SC mirror commonly observed functional links between the anterior hippocampus and temporal regions and between the posterior hippocampus and parietal/occipital regions (16, 40–43). Our results provide new insights into the neuroanatomical architecture of these functional associations in the human brain. Further interpretation of these observations are facilitated by the results of our more detailed endpoint density analyses and are discussed below.

### Endpoint density mapping of human cortico-hippocampal connectivity

To date, our knowledge of human cortico-hippocampal anatomical connectivity is largely inferred from the results of tract-tracing studies conducted in non-human primates and rodents. We know much less about these patterns in the human brain. To address this gap, we adapted a tractography pipeline to track streamlines entering/leaving the hippocampus, identify the location of their ‘endpoints’ and create spatial distribution maps of endpoint density within the hippocampus associated with each cortical area. The resulting EDMs allowed us to quantitatively assess and visualise where, within the hippocampus, different cortical areas preferentially ‘connect’. To our knowledge, this is the first time such a specific approach has been used to map anatomical connectivity of the human hippocampus *in-vivo*.

We observed striking differences in the location of endpoint density along both the anterior-posterior and medial-lateral axes of the hippocampus. For example, temporopolar and inferolateral temporal cortical areas displayed the greatest endpoint density along the lateral aspect of the head and body of the hippocampus (Supplementary Figure S4a-e; aligning with the location of distal CA1/proximal subiculum; see Figure 5a) and in a circumscribed cluster in the anterior medial hippocampus at the level of the uncal apex (Supplementary Figure S4a-d; aligning with the location of distal subiculum/proximal presubiculum; see Figure 5a). In contrast, areas in the medial parietal and occipital cortices displayed the greatest endpoint density along the medial aspect of the tail and body of the hippocampus (Supplementary Figures S3a-e (medial parietal) and 5a-e (occipital); aligning with the location of the distal subiculum/proximal presubiculum; see Figure 5a). We discuss these observations in relation to the non-human primate literature below.

### Comparison to tract-tracing data in non-human primates

The results of our endpoint density analyses broadly overlap with observations from the non-human primate literature. For simplicity, we demonstrate this by comparing the results for three cortical areas with equivalent observations from the series of tract-tracing investigations of the macaque hippocampus described by Insausti & Muñoz (22). In the macaque, retrograde tracer injection into the RSC results in localised labelling of cells in the posterior presubiculum. Our results revealed high endpoint densities associated with the RSC were primarily localised to a homologous region in the posterior medial hippocampus (distal subiculum/proximal presubiculum; brown ellipsoids in Figure 5b). In the macaque, retrograde tracer injection into the PeEc results in dense labelling of cells along the anterior-posterior axis of the hippocampus primarily localised to the CA1/subiculum transition area and the presubiculum. Our results mirrored this pattern. We observed areas of endpoint density along the lateral hippocampus (blue ellipsoids in Figure 5c; distal CA1/proximal subiculum) and along the medial hippocampus (brown ellipsoids in Figure 5c; distal subiculum/proximal presubiculum). In the macaque, labelling associated with area TE is scarce compared to that of the PeEc with labelled cells localised to the posterior CA1/subiculum transition area and the posterior presubiculum. Again, our results broadly aligned with this pattern. Compared with PeEc, we observed less endpoint density associated with TE2a (roughly corresponding to the injection site described by Insausti & Muñoz) which was primarily localised to the lateral (blue ellipsoids in Figure 5d; distal CA1/proximal subiculum) and medial (brown ellipsoids in Figure 5d; distal subiculum/proximal presubiculum) hippocampus. In contrast to the macaque, however, endpoint density was most strongly expressed in the lateral aspect of the anterior hippocampus (red ellipsoid in Figure 5d). We discuss potential interpretations for this and other differences later in the discussion. Overall, our results broadly aligned with patterns observed in the non-human primate brain and provide new and detailed insights regarding where specific cortical areas preferentially connect within the human hippocampus.

### New evidence for hubs of anatomical connectivity in the human hippocampus?

High endpoint density within the hippocampus was restricted to areas that aligned with the location of CA1, subiculum and the pre- and parasubiculum and was notably absent from areas aligning with DG/CA4 (hilus), CA3 and CA2. These observations mirror reports in the rodent and non-human primate literature where non-EC cortical areas predominantly connect with subicular cortices and the CA1/subiculum transition area (20, 22). Indeed, in non-human primates, the CA1/subiculum transition area and the presubiculum appear to be ‘hotspots’ of anatomical connectivity for multiple cortical areas (22, 44). In accordance with this, we observed that these specific regions within the human hippocampus displayed high endpoint density for multiple cortical areas. For example, the anterior lateral hippocampus (aligning with the distal CA1/proximal subiculum) displayed high endpoint density for multiple temporal cortical areas (Figure 3a) and the posterior medial hippocampus (aligning with the distal subiculum/proximal presubiculum) displayed high endpoint density for multiple areas of the medial parietal and occipital cortices (Figure 3b and c respectively). Another cluster of high endpoint density in the anterior medial hippocampus (aligning with the distal subiculum/proximal presubiculum) was more broadly associated with temporal, medial parietal and occipital areas (Figure 3a-d). Taken together, these observations provide evidence that discrete hubs of dense anatomical connectivity may exist along the anterior-posterior axis of the human hippocampus and that these hubs align with the location of the CA1/subiculum transition area and the distal subiculum/proximal presubiculum.

To the best of our knowledge, this is the first quantitative report of these patterns in the human brain and we highlight the intriguing possibility that two medial hippocampal hubs of high anatomical connectivity may exist; a posterior medial hub preferentially linked with visuospatial processing areas in medial parietal and occipital cortices and an anterior medial hub more broadly linked with temporopolar, inferotemporal, medial parietal and occipital areas. We tentatively speculate that these circumscribed areas of the medial hippocampus could represent highly connected hubs of information flow between the hippocampus and distributed cortical networks. Indeed, the cortical areas identified in this study may represent key areas for direct cortico-hippocampal interactions in support of episodic/semantic memory processing and consolidation.

Accumulating evidence from neuroimaging and clinical studies suggest that the medial hippocampus plays an important role in visuospatial cognition. The anterior medial hippocampus is consistently engaged during cognitive tasks that require processing of naturalistic scene stimuli in aid of episodic memory (45), visuospatial mental imagery (5), perception and mental construction (e.g., imagination) of scenes (46, 47). Strikingly, the location of the anterior medial cluster of anatomical connectivity observed in the current study aligns with the location of functional clusters commonly observed in functional MRI investigations of these cognitive processes (5, 45–48) (see Figure 6 for comparison). Furthermore, recent evidence from the clinical domain suggests that the posterior medial hippocampus may be a critical hub in a broader memory circuit (49). The medial hippocampus has more broadly been proposed as a putative hippocampal hub for visuospatial cognition (50). Our results lend further support to these proposals by showing that specific regions of both the anterior and posterior medial hippocampus display dense anatomical connectivity with multiple cortical areas in the human brain.

**Figure 6.**
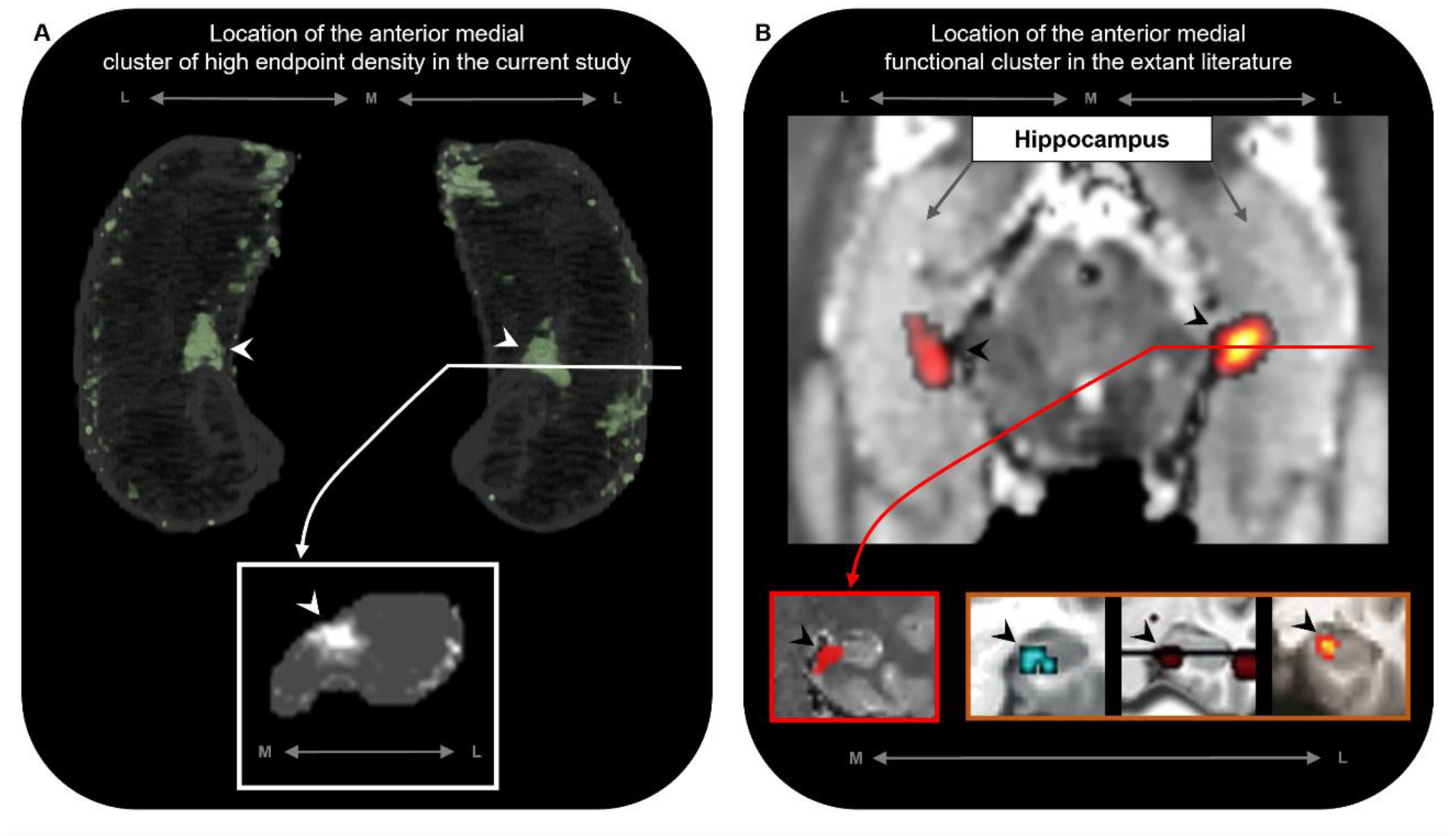
The location of the anterior medial anatomical cluster aligns with the location of a commonly observed anterior medial functional cluster. a. A 3D rendered representation of the bilateral group level hippocampus mask (top; transparent grey) is presented overlaid with the endpoint density map averaged across the most highly connected brain areas in temporal, medial parietal and occipital cortices (green; see Figure 3d for details); a representative slice of hippocampus in the coronal plane (bottom panel; grey) at the level of the uncal apex (indicated by white line) is presented overlaid with the endpoint density map (white). b. An axial section of a T2-weighted image (top; from a separate study) showing the bilateral hippocampus overlaid with the location of a circumscribed functional cluster observed in the anterior medial hippocampus during a functional MRI investigation of visuospatial mental imagery (5). A representative slice of the hippocampus (bottom left panel; red border) at the level of the uncal apex (indicated by red line) is presented to show the location of this anterior medial functional cluster in the coronal plane (images reproduced with permission from (5)). Circumscribed functional clusters in the anterior medial hippocampus are commonly observed in studies of ‘scene-based visuospatial cognition’ such as episodic memory and prospection (bottom right panel; orange border). Images reproduced with permission from (46) (left), (45) (middle) and (47) (right). Note, the location of these commonly observed functional clusters in the anterior medial hippocampus (black arrows in panel b) align with the location of the anatomical cluster in the anterior medial hippocampus observed in the current study (white arrows in panel a). M=medial; L=lateral.

In addition, our results provide new anatomical insights to inform current debates on functional differentiation along the anterior-posterior axis of the human hippocampus. Our anterior-posterior axis analyses revealed that specific cortical areas displayed a gradient style increase in connectivity strength along the anterior-posterior axis of the hippocampus while others displayed non-linear patterns of connectivity. In parallel, the results of our endpoint density analyses suggest that discrete clusters of dense connectivity may also exist within the hippocampus. Together, these observations provide new evidence to support recent proposals that both gradients and circumscribed parcels of extrinsic connectivity may exist along the anterior-posterior axis of the human hippocampus (13–15) and that different circuits may exist within the hippocampus, each associated with different cortical inputs that underpin specific cognitive functions (5). How the complex patterns of anatomical connectivity observed in the current study relate to functional differentiation along the long-axis of the hippocampus (11–14) and it’s subfields (16, 17) will be a fruitful area of research in coming years.

### Are there human specific patterns of cortico-hippocampal connectivity?

Despite areas of concordance with the non-human primate literature noted above, we also observed important differences. As noted earlier, we observed broader patterns of endpoint density for area TE2a than we expected based on the non-human primate literature. Also, non-human primate studies have found direct and substantial connectivity between the hippocampus and orbitofrontal and superior temporal cortices (22). We found only weak patterns of connectivity between the hippocampus and these regions. Also of note, in contrast to the non-human primate literature, we observed dense patterns of anatomical connectivity between the posterior medial hippocampus and early visual processing areas in the occipital lobe. It is important to note that our methods differ significantly from the carefully controlled injection of tracer into circumscribed portions of the brain. MRI investigations of SC are inherently less precise, and methodological limitations likely explain some of our observed differences. It is important to note, however, that while we expect to see evolutionarily conserved overlaps in SC between macaque and human brains, we should not expect exactly the same patterns of connectivity. Since splitting from a common ancestor, macaques and humans have likely evolved species specific patterns of cortico-hippocampal connectivity to support species specific cognitive functions. Whether differences observed in the current study reflect methodological limitations, species differences or perhaps most likely, a mix of both, require further investigation.

To our knowledge, only one prior study has attempted to characterise the broader hippocampal ‘connectome’ in the healthy human brain (38). Our study differs from this recent report in several important technical aspects. We analysed connectivity profiles of the head, body and tail portions of the hippocampus separately in addition to the whole hippocampus. We manually delineated the hippocampus for each participant to ensure full coverage of the hippocampus and to minimise the hippocampus mask ‘spilling’ into adjacent white matter (a common occurrence with automated segmentation methods; see Supplementary Figure S6). We used the HCPMMP scheme, which provides more detailed subdivisions of cortical grey matter (39). We also included SIFT2 (51) in our analysis pipeline to increase biological accuracy of quantitative connectivity estimates. Finally, we used a carefully adapted tractography method that incorporates anatomical constraints and allowed streamlines to enter/leave the hippocampus, which provided us with a means to determine the location and topography of streamline ‘endpoints’, and therefore their distribution within the hippocampus (see Methods for detail relating to each of these points).

In closing, this study represents a first attempt to apply this method and has some limitations. Specifically, our method relies on several manual steps to delineate the hippocampus and to amend the grey matter-white matter interface that abuts the inferior portion of the hippocampus (described in Materials and Methods). This can be time consuming and requires expertise to accurately identify the hippocampus along its entire anterior-posterior extent on structural MRI scans. However, considering automated methods of hippocampal segmentation are sometimes not sufficiently accurate, particularly in the anterior and posterior most extents of the hippocampus (see Supplementary Figure S6 for representative examples), manual delineation is the gold standard and ensures the best results. We restricted our analysis to a limited number of subjects under the age of 35 and selected participants whose hippocampus was clearly visible along its entire anterior-posterior axis on T1-weighted structural scans. While this ensured we used the best data quality available, further work should explore how these results may differ in the context of healthy ageing and in diseases that affect the hippocampus such as Alzheimers disease, epilepsy and schizophrenia. How reliable our pipeline is in data acquired with more traditional clinical protocols remains to be explored and was beyond the scope of the current study. Despite the high-quality HCP data used in this study, limitations in spatial resolution likely restrict our ability to track particularly convoluted white-matter pathways within the hippocampus and our results should be interpreted with this in mind. Future work using higher resolution data will allow more targeted investigations and is necessary to confirm or refute the patterns we observed here. These limitations notwithstanding, our results provide new detailed insights into anatomical connectivity of the human hippocampus, can inform theoretical models of human hippocampal function as they relate to the long-axis of the hippocampus and can help fine-tune network connectivity models.

From a clinical perspective, the hippocampus is central to several neurodegenerative and neuropsychiatric disorders. Considering healthy memory function is dependent upon the integrity of white-matter fibres that connect the hippocampus with the rest of the brain, developing a more detailed understanding of the anatomical connectivity of the anterior-posterior axis of the hippocampus and its subfields has downstream potential to help us better understand hippocampal-dependent memory decline in ageing and clinical populations. Our novel method can potentially be harnessed to measure changes in anatomical connectivity between the hippocampus and cortical areas known to be affected in neurodegenerative diseases, to assist with monitoring of disease progression and/or as a diagnostic tool. In addition, our method could be deployed to visualise patient specific patterns of hippocampal connectivity to support surgical planning in patients who require MTL resection for intractable MTL epilepsy.

## Materials and Methods

### Participant details

Ten subjects (7 female) were selected from the minimally processed Human Connectome Project (HCP) 100 unrelated subject database (<35yo). Subjects were selected based on the scan quality and visibility of the outer boundaries of the hippocampus on each participants T1-weighted structural MRI scan. This was done in order to increase the anatomical accuracy of our hippocampal segmentations (described below).

### Image acquisition

The HCP diffusion protocol consisted of three diffusion-weighted shells (b-values: 1000, 2000 and 3000 s/mm2, with 90 diffusion weighting directions in each shell) plus 18 reference volumes (b = 0 s/mm2). Each diffusion-weighted image was acquired twice, with opposite phase-encoded direction to correct for image distortion (52). The diffusion image matrix was 145 × 145 with 174 slices and an isotropic voxel size of 1.25 mm. The TR and TE were 5520 and 89.5 ms, respectively. Each subject also included a high resolution T1-weighted dataset, which was acquired with an isotropic voxel size of 0.7 mm, TR/TE = 2400/2.14 ms, and flip angle = 8°.

### Manual segmentation of the hippocampus

The whole hippocampus was manually segmented for each participant on coronal slices of the T1-weighted image using ITK-SNAP (53). Although automated methods of hippocampal segmentation are available, they are sometimes not sufficiently accurate particularly in the anterior and posterior most extents of the hippocampus (see Supplementary Figure S6 for representative examples). Although labour intensive, manual segmentation by an expert in hippocampal anatomy remains the gold-standard for detailed and accurate investigation of the human hippocampus. We adapted the manual segmentation protocol outlined by Dalton and colleagues (54). While this protocol details a method for segmenting hippocampal subfields, we followed guidelines as they relate to the outer boundaries of the hippocampus. This ensured that the whole hippocampus mask for each participant contained all hippocampal subfields (dentate gyrus, CA4-1, subiculum, presubiculum and parasubiculum) and encompassed the entire anterior-posterior extent of the hippocampus (see (54) for details). Representative examples of the hippocampus mask are presented in Supplementary Figures S6 and S7. Manual segmentations were conducted by an expert in human hippocampal anatomy and MRI investigation of the human hippocampus (M. A. D) with 14 years experience including histological (55) and MRI investigations (5, 16, 17, 54). For the anterior-posterior axis analysis, we split each participants whole hippocampus mask into thirds corresponding with the head, body and tail of the hippocampus. This was done in accordance with commonly used anatomical landmark-based methods. In brief, the demarcation point between the head and body of the hippocampus was the uncal apex (10, 56) and the demarcation point between the body and tail of the hippocampus was the anterior most slice in which the crus of the fornix was fully visible (57, 58).

### Image pre-processing and whole-brain tractography

Besides the steps carried out by the HCP team as part of the minimally processed datasets, the additional image processing pipeline included in our analysis is summarised in Supplementary Figure S8a. Processing was performed using the MRtrix software package (http://www.mrtrix.org) (59, 60). Additional processing steps were implemented in accordance with previous work (61) and included bias-field correction (62) as well as multi-shell multi-tissue constrained spherical deconvolution to generate a fibre orientation distribution (FOD) image (63–65). The T1 image was used to generate a ‘five-tissue-type’ (5TT) image using FSL (66–70); tissue 1 = cortical grey matter, tissue 2 = sub-cortical grey matter, tissue 3 = white matter, tissue 4 = CSF, and tissue 5 = pathological tissue. The FOD image and the 5TT image were used to generate 70 million anatomically constrained tracks (66) using dynamic seeding (71) and the 2nd-order Integration over Fibre Orientation Distributions (iFOD2 (72)) probabilistic fibre-tracking algorithm. The relevant parameters included: 70 million tracks, dynamic seeding, backtracking option specified, FOD cut-off 0.06, minimum track length 5 mm, maximum track length 300 mm, maximum of 1000 attempts per seed.

### Hippocampus tractography

We developed a tailored pipeline to track streamlines into the hippocampus. To do this, we first amended the gm-wm interface immediately inferior to the hippocampus. This was necessary because the manually segmented hippocampus mask lay slightly superior to the automatically generated gm-wm interface (see Supplementary Figure S7b; middle image). Pilot testing showed that streamlines terminated when reaching this portion of the gm-wm interface, thereby impeding streamlines from traversing the inferior border of the hippocampus. This was a problem because white matter fibres innervate the hippocampus primarily through this region (and also via the fimbria/fornix) (73). It was, therefore, important to ensure that streamlines could cross the inferior border of the hippocampus mask in a biologically plausible manner. To facilitate this, we created an additional hippocampus mask for each participant that extended inferiorly to encompass portions of the gm-wm border that lay immediately inferior to the hippocampus (see Supplementary Figure S7b; right image). This amended hippocampus mask was labelled as white matter in the modified 5TT image (referred to as m5TT). This served to remove the portion of the gm-wm border immediately inferior to the hippocampus and ensured that streamlines could enter/leave the hippocampus in a biologically plausible manner. Additionally, the original whole hippocampus segmentation was assigned as 5th tissue type in the m5TT image (i.e. where no anatomical priors are applied within the ACT framework in MRtrix; Supplementary Figure S8b). This allowed streamlines to move within the hippocampus.

In summary, amending the erroneous gm-wm border allowed streamlines to traverse hippocampal boundaries in a biologically plausible manner and labelling the manually segmented hippocampus as a 5th tissue type permitted streamlines to move within the hippocampus. Together, this allowed us to follow the course of each streamline within the hippocampus and determine the location of each streamline “endpoint” (described below).

Next, the FOD image was used with the m5TT image to generate an additional 10 million tracks. This set of anatomically constrained tracks (66) were seeded from the manually segmented hippocampus, and iFOD2 was used for fibre tracking(72). The 70 million whole-brain tracks and the 10 million hippocampus tracks were combined, and spherical-deconvolution informed filtering of tractograms 2 (SIFT2 (71)) was used on the combined 80 million track file, thereby assigning a weight to each track and providing biological credence to the connectivity measurements (74). Within the SIFT2 framework, connectivity is then computed not by counting the number of tracks but by the sum of its SIFT2 weights. Tracks (and SIFT2 weights) which had an endpoint in the hippocampus were extracted (referred to here as the ‘hippocampus tractogram’) and used in both the whole hippocampus and anterior-posterior axis analyses.

### Whole hippocampus connectivity

FreeSurfer (75) was used to further process the T1-weighted image. The Human Connectome Project Multi-Modal Parcellation 1.0 (HCPMMP) (39) was mapped to each subject in accordance with previous work (76). The parcellation divided the cerebral cortex into 360 parcels (180 per hemisphere). Importantly, we replaced the automated hippocampus and presubiculum parcels with the manually segmented hippocampus (which included the presubiculum) for greater anatomical accuracy (referred to as ‘modified HCPMMP’; Supplementary Figure S8c). The structural connectivity of tracks between the hippocampus and the other parcels was obtained using tracks (and SIFT2 weights) from the hippocampus and the modified HCPMMP (containing the whole-hippocampus segmentation). The strength of connectivity between the hippocampus and every other parcel of the HCPMMP was measured by the sum of the SIFT2 weighted connectivity values (51). For each parcel (cortical area), we combined left and right hemisphere values (i.e., left and right retrosplenial cortex) and report bilateral results.

### Anterior-posterior axis connectivity

As described in the manual segmentation section, the whole hippocampus was subsequently divided into thirds (head, body and tail, as shown in Supplementary Figure S8d). For the anterior-posterior axis analyses, each of these three regions were added to the HCPMMP as their own unique parcel. In a similar manner to the whole hippocampus connectivity analysis, the strength of connectivity between each hippocampal region (head, body and tail) and each parcel of the modified HCPMMP was measured by the sum of the SIFT2 weighted connectivity values. To assess whether connectivity values between each cortical brain area and the head, body and tail portions of the hippocampus were statistically significant, we conducted Bonferroni-corrected paired-samples t-tests. These are reported in the main text when significant at a level of p < 0.05.

### Track Density Image (TDI) maps and endpoint creation

#### TDI mapping of tracks between the whole-hippocampus and the other parcels

The extracted hippocampus tractogram was used to isolate tracks (and weights) between the whole hippocampus parcel and every other parcel in the modified HCPMMP file. Two different TDI maps (77, 78) were computed for each parcel; a TDI of the hippocampus tractogram, and a TDI map showing only the endpoints of this tractogram. Both TDI maps were constructed at 0.2 mm isotropic resolution. These TDI endpoint maps were used in the group-level analysis described below. Note, we refer to these TDI endpoint maps as ‘endpoint density maps’ (EDMs) in the main text.

### Group level analysis of TDI endpoint maps

#### Group level hippocampus template and TDI endpoint map registration

We employed the symmetric group-wise normalization method (SyGN) (79) implemented in the ANTs toolbox (http://stnava.github.io/ANTs/) to build a population-specific hippocampus template. Specifically, the cross-correlation metric was used to optimize the boundary agreement among the hippocampi masks of each participant. Then, each individual hippocampus mask was registered to the generated population template with the combined linear and non-linear transformation. For each participant, the transformation parameters were recorded and applied to the TDI endpoint maps which were warped into the template space at a resolution of 0.7 mm isotropic. The group average was then calculated providing a group-level distribution map of endpoint density within the hippocampus for each parcel of interest. EDMs were visualised in mrview (the MRtrix image viewer). Representative images displayed in our figures were visualised with the minimum and maximum intensity scale set at 0 and 0.05 respectively and a minimum and maximum threshold set at 0.02 and 0.5 respectively.

## Data availability

We used data from the Human Connectome Project (HCP) 100 unrelated subject database; https://www.humanconnectome.org/study/hcp-young-adult/data-releases. Requests for further information regarding our analysis pipeline should be directed to the corresponding author, Marshall Dalton.

## Author Contributions

M.A.D. and F.C. conceived the experiment. M.A.D., A.D. and J.L. analyzed the data. M.A.D. wrote the manuscript with input from F.C., A.D. and J.L.

## Competing Interest Statement

The authors declare no competing interests.

## Acknowledgments

Data were provided by the Human Connectome Project, WU-Minn Consortium (Principal Investigators: David Van Essen and Kamil Ugurbil; 1U54MH091657) funded by the 16 NIH Institutes and Centers that support the NIH Blueprint for Neuroscience Research; and by the McDonnell Center for Systems Neuroscience at Washington University, St. Louis, MO.

We are grateful for the support of the National Health and Medical Research Council of Australia (grant numbers APP1091593 and APP1117724), and the Australian Research Council (grant number DP170101815).

The authors acknowledge the technical assistance provided by the Sydney Informatics Hub and Sydney Imaging, two Core Research Facilities of the University of Sydney, Australia.

## Supplementary Information

Note S1: Additional medial temporal lobe analyses.

While the primary focus of our study was to characterise anatomical connectivity between the hippocampus and non-medial temporal lobe (MTL) cortical areas, we also characterised patterns of connectivity between the hippocampus and MTL cortical areas. Cortical areas within the MTL were highly connected with the hippocampus and cumulatively accounted for 52% of all cortical connections. The most highly connected area was the EC (24% of all cortical connections) followed by PeEc (14%), PHA2 (6%), PHA1 (5%) and PHA3 (3%) (see Figure S1a and Table S1). Results of the anterior-posterior axis analyses revealed that the EC and PeEc displayed a gradient style anterior-to-posterior decrease in connectivity. In contrast, PHA1-3 each displayed the highest degree of connectivity with the body of the hippocampus and lower connectivity with both the head and tail of the hippocampus (Figure S1b and Table S2).

The results of the group level endpoint analyses showed that MTL cortical areas had dense patterns of connectivity with the hippocampus. For example, the EC displayed high endpoint density along the entire anterior-posterior axis of the hippocampus. Although difficult to visualise due to the high density of endpoints, the results of our anterior-posterior analysis showed that endpoint density was greatest in the head and body of the hippocampus (see Figure S1b and Table S2). Endpoint density was primarily located in portions of the hippocampus aligning with the location of the CA1, subiculum, pre- and parasubiculum (See Figure S2a).

Endpoint density expression associated with PeEc differed along the anterior-posterior axis of the hippocampus. In the hippocampal head, endpoint density was more pronounced in the lateral hippocampus primarily aligning with the location of the CA1 and subiculum (indicated by blue arrows in Figure S2b). Moving into the body, expression in the CA1 and subiculum was maintained (indicated by blue arrows in Figure S2b) and additional clusters of endpoint density were expressed in the medial hippocampus aligning with the location of the distal subiculum/proximal presubiculum (indicated by white arrows in Figure S2b). Moving into the hippocampal tail, clusters of endpoint density appeared to be more localised to a lateral region of the hippocampus aligning with the location of distal CA1/proximal subiculum (indicated by blue arrows in Figure S2a) with comparatively weaker expression in medial portions of the hippocampal tail.

Endpoint density associated with PHA1-3 was most pronounced in the body of the hippocampus and primarily localised to lateral areas aligning with the location of CA1 and subiculum (indicated by blue arrows in Figure S2c-e) and in the medial hippocampus aligning with the location of the distal subiculum/proximal presubiculum (indicated by white arrows in Figure S2c-e). PHA1-3 expressed less endpoint density in the hippocampal head which was localised to the lateral hippocampus aligning with the location of CA1 and subiculum (indicated by blue arrows in Figure S2c-e). Moving into the hippocampal tail, clusters of endpoint density were more localised to a lateral region aligning with the location of distal CA1/proximal subiculum (indicated by blue arrows in Figure S2c-e). However, PHA1-3 showed different patterns of endpoint density in the medial aspect of the hippocampal tail. PHA1 and, to a lesser extent, PHA2 displayed endpoint density in the posterior medial hippocampus aligning with the location of the distal subiculum/proximal presubiculum (indicated by yellow arrows in Figure S2c and d). In contrast, PHA3 displayed modest endpoint density in the posterior medial hippocampus.

---

**Fig. S1.**
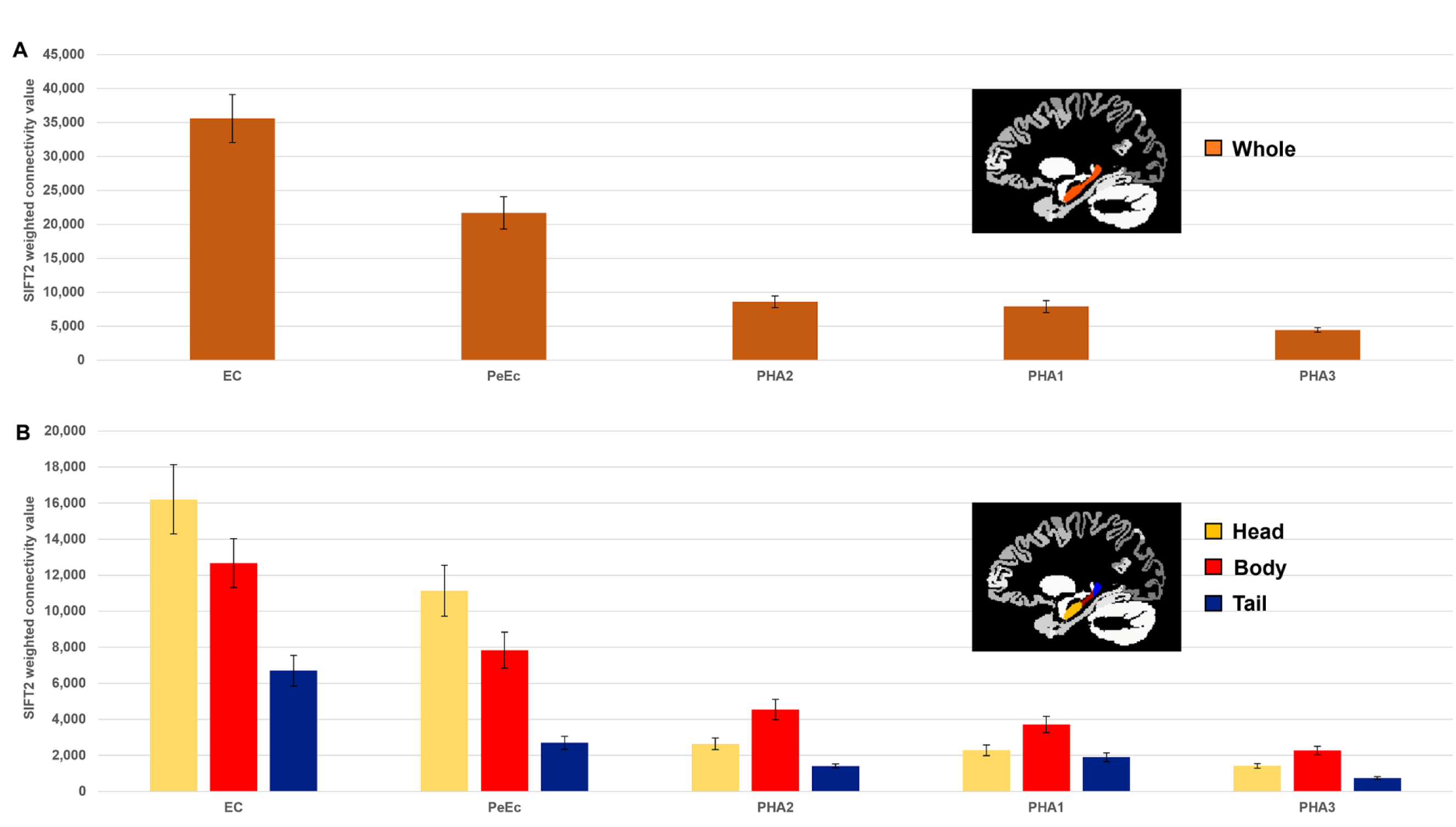
MTL cortices anatomical connectivity with the hippocampus. a. Histogram plotting the mean structural connectivity (given by the sum of SIFT2 weighted values) associated with MTL cortical areas connected with the whole hippocampus. Error bars represent the standard error of the mean. b. Histogram plotting the corresponding mean SIFT2 weighted values associated with anterior (yellow), body (red) and tail (blue) portions of the hippocampus for MTL cortical areas presented in a. Errors bars represent the standard error of the mean.

**Fig. S2.**
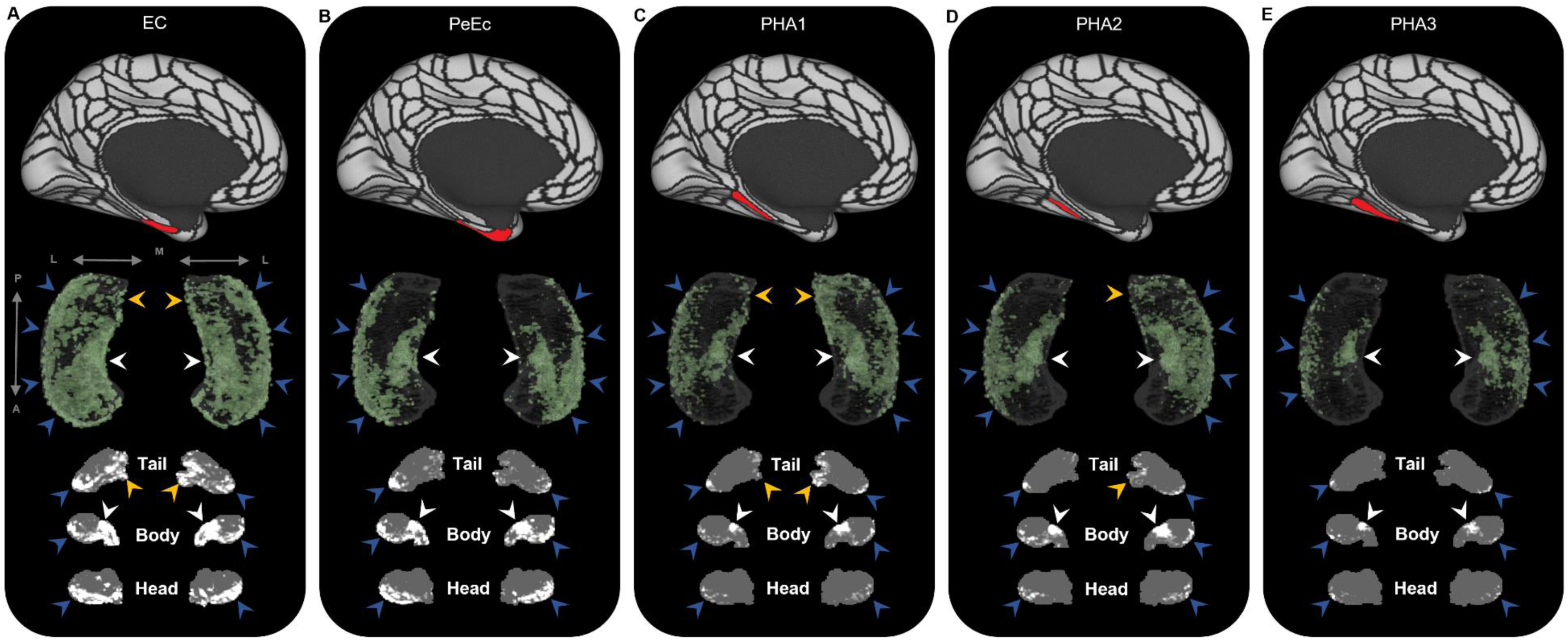
Representative examples of the spatial distribution of endpoint density within the hippocampus for MTL brain areas. Representative examples of the location of endpoint densities associated with EC (a), PeEc (b) PHA1 (c), PHA2 (d) and PHA3 (e). In each panel, the location of the relevant brain area is indicated in red on the brain map (top); a 3D rendered representation of the bilateral group level hippocampus mask (middle; transparent grey) is presented overlaid with the endpoint density map associated with each brain area (green); representative slices of the head, body and tail of the hippocampus are displayed in the coronal plane (bottom; grey) and overlaid with endpoint density maps (white). Note, as expected, the spatial distribution of endpoint density associated with MTL cortical areas is denser than that observed for non-MTL cortical areas (compare with Figures S3-5). EC displayed high endpoint density throughout the hippocampus (a). PeEc (b) and PHA3 (e) displayed the greatest endpoint density along the anterior-posterior extent of the lateral hippocampus (blue arrows) and in a circumscribed region in the anterior medial hippocampus (white arrows). PeEc and PHA3 showed little endpoint density in the posterior medial hippocampus. PHA1 (c) and PHA2 (d) displayed high endpoint density along the anterior-posterior extent of the lateral hippocampus (blue arrows), in the anterior medial hippocampus (white arrows) and, in contrast to PeEc and PHA3, in the posterior medial hippocampus (yellow arrows). A=anterior; P=posterior; M=medial; L=lateral.

**Fig. S3.**
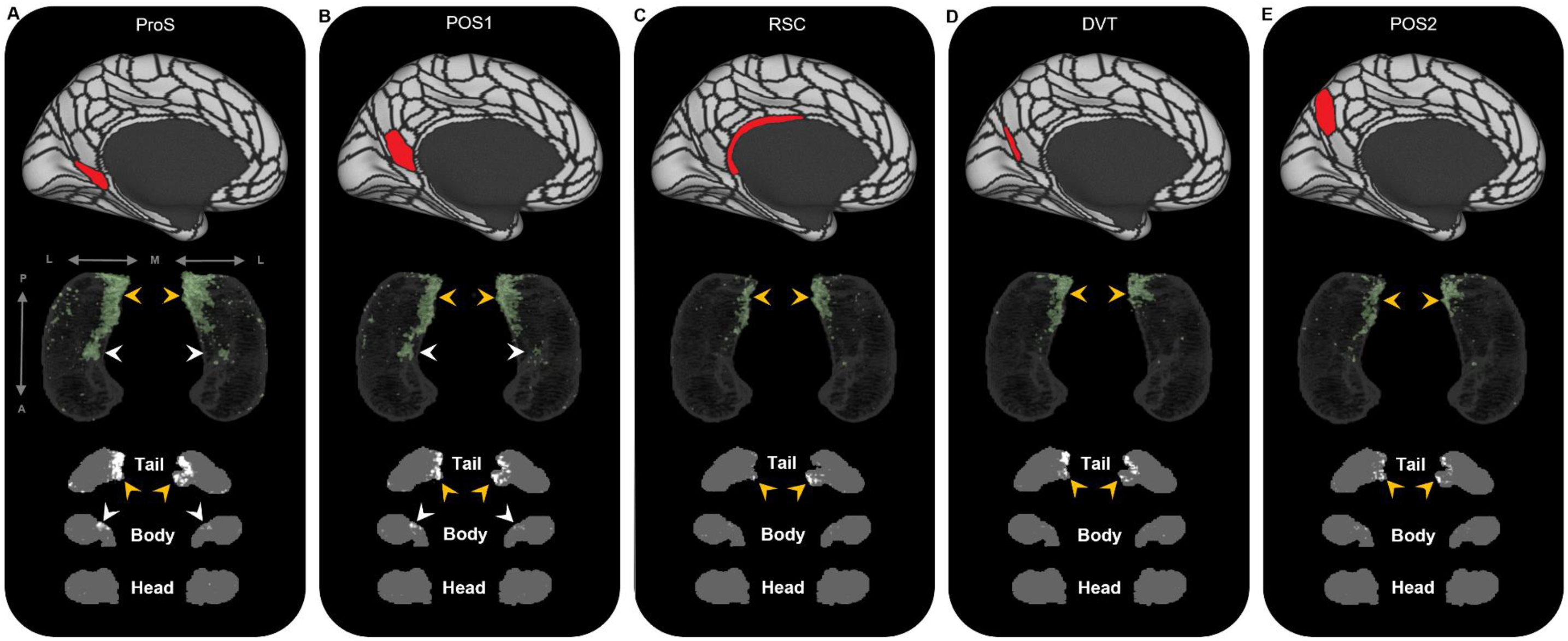
Representative examples of the spatial distribution of endpoint density within the hippocampus associated with the five most highly connected medial parietal brain areas. Representative examples of the location of endpoint densities associated with ProS (a), POS1 (b) RSC (c), DVT (d) and POS2 (e). In each panel, the location of the relevant brain area is indicated in red on the brain map (top); a 3D rendered representation of the bilateral group level hippocampus mask (middle; transparent grey) is presented overlaid with the endpoint density map associated with each brain area (green); representative slices of the head, body and tail of the hippocampus are displayed in the coronal plane (bottom; grey) and overlaid with endpoint density maps (white). Note, the spatial distribution of endpoint density within the hippocampus associated with each of these medial parietal brain areas is primarily localised to the posterior medial hippocampus (yellow arrows in panels a-e). ProS and POS1 also display clusters of endpoint density in a circumscribed region in the anterior medial hippocampus (white arrows in panels a and b). A=anterior; P=posterior; M=medial; L=lateral.

**Fig. S4.**
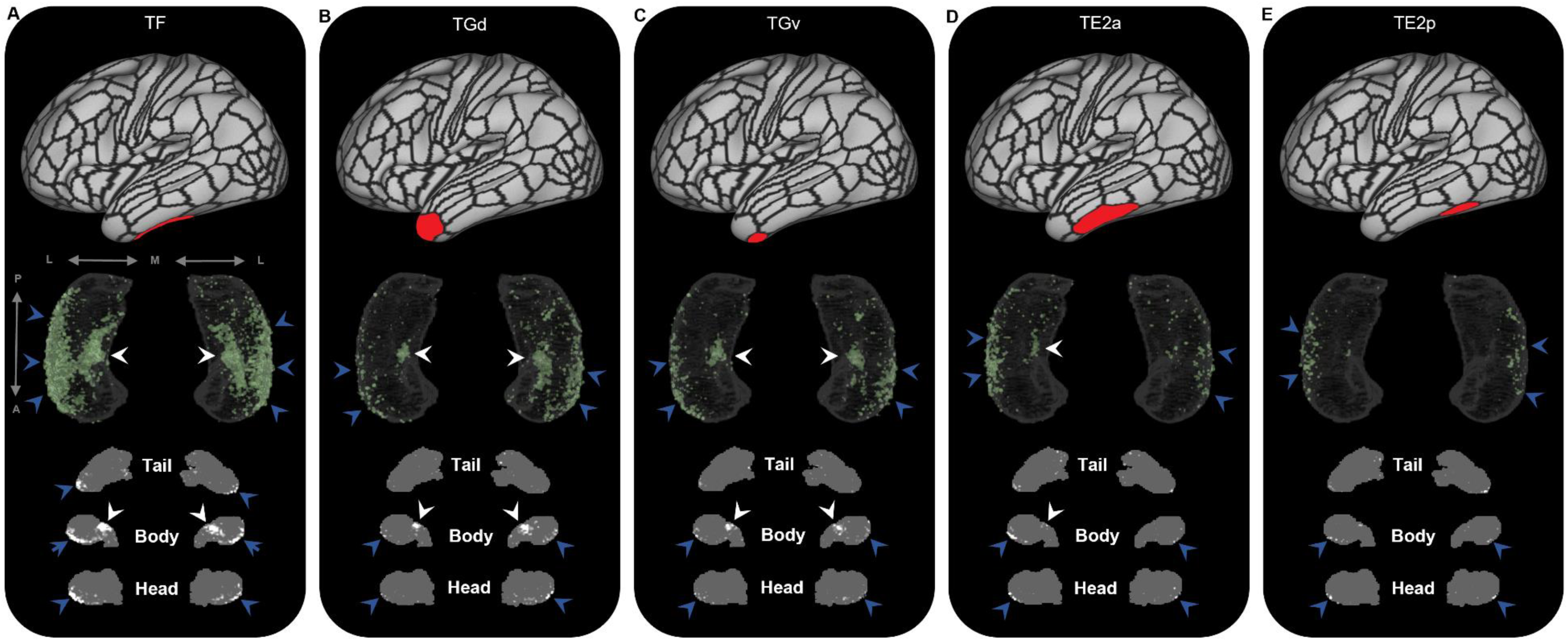
Representative examples of the spatial distribution of endpoint density within the hippocampus associated with the five most highly connected non-MTL temporal brain areas. Representative examples of the location of endpoint densities associated with TF (a), TGd (b) TGv (c), TE2a (d) and TE2p (e). In each panel, the location of the relevant brain area is indicated in red on the brain map (top); a 3D rendered representation of the bilateral group level hippocampus mask (middle; transparent grey) is presented overlaid with the endpoint density map associated with each brain area (green); representative slices of the head, body and tail of the hippocampus are displayed in the coronal plane (bottom; grey) and overlaid with endpoint density maps (white). Note, the spatial distribution of endpoint density within the hippocampus associated with each of these temporal brain areas is primarily localised to portions of the lateral hippocampal head and body (blue arrows in panels a-e) and a circumscribed region in the anterior medial hippocampus (white arrows in panels a-d). A=anterior; P=posterior; M=medial; L=lateral.

**Fig. S5.**
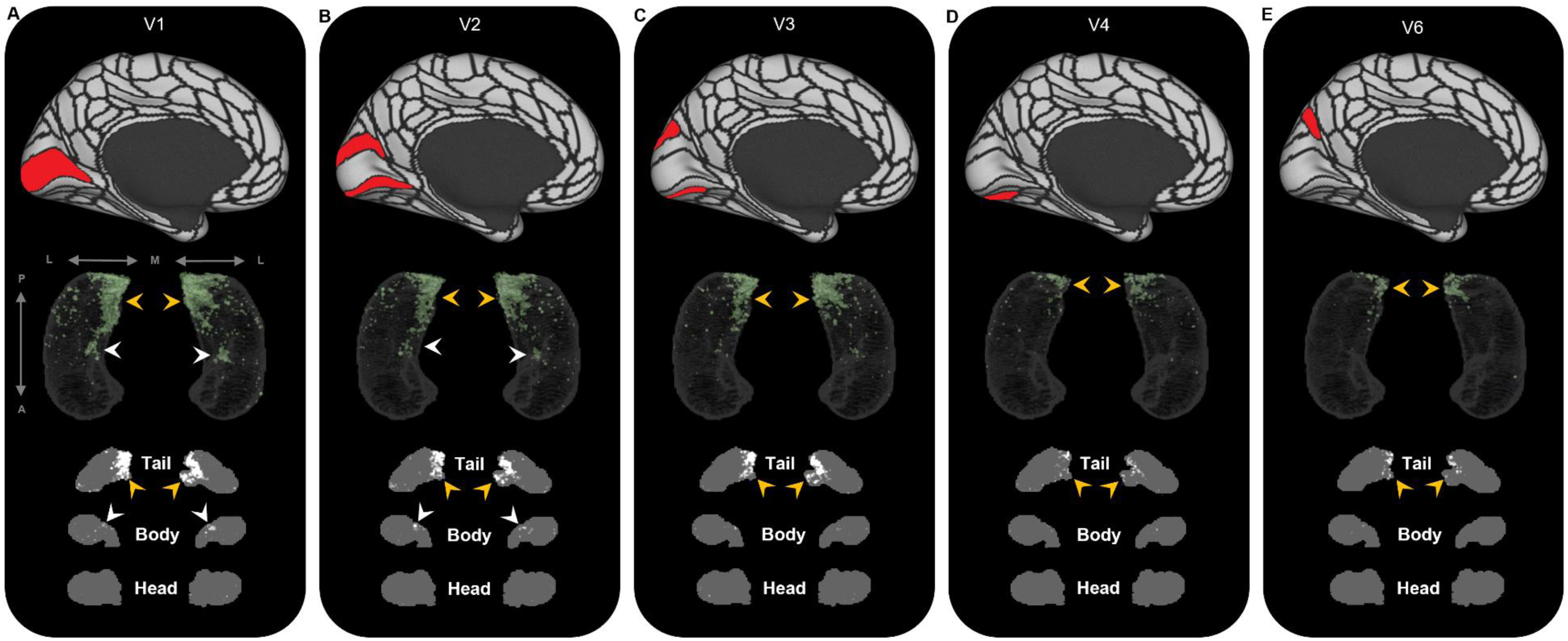
Representative examples of the spatial distribution of endpoint density within the hippocampus associated with the five most highly connected occipital brain areas. Representative examples of the location of endpoint densities associated with V1 (a), V2 (b) V3 (c), V4 (d) and V6 (e). In each panel, the location of the relevant brain area is indicated in red on the brain map (top); a 3D rendered representation of the bilateral group level hippocampus mask (middle; transparent grey) is presented overlaid with the endpoint density map associated with each brain area (green); representative slices of the head, body and tail of the hippocampus are displayed in the coronal plane (bottom; grey) and overlaid with endpoint density maps (white). Note, the spatial distribution of endpoint density within the hippocampus associated with each of these occipital brain areas is primarily localised to the posterior medial hippocampus (yellow arrows in panels a-f). V1 and V2 also display clusters of endpoint density in a circumscribed region in the anterior medial hippocampus (white arrows in panels a and b). A=anterior; P=posterior; M=medial; L=lateral.

**Fig. S6.**
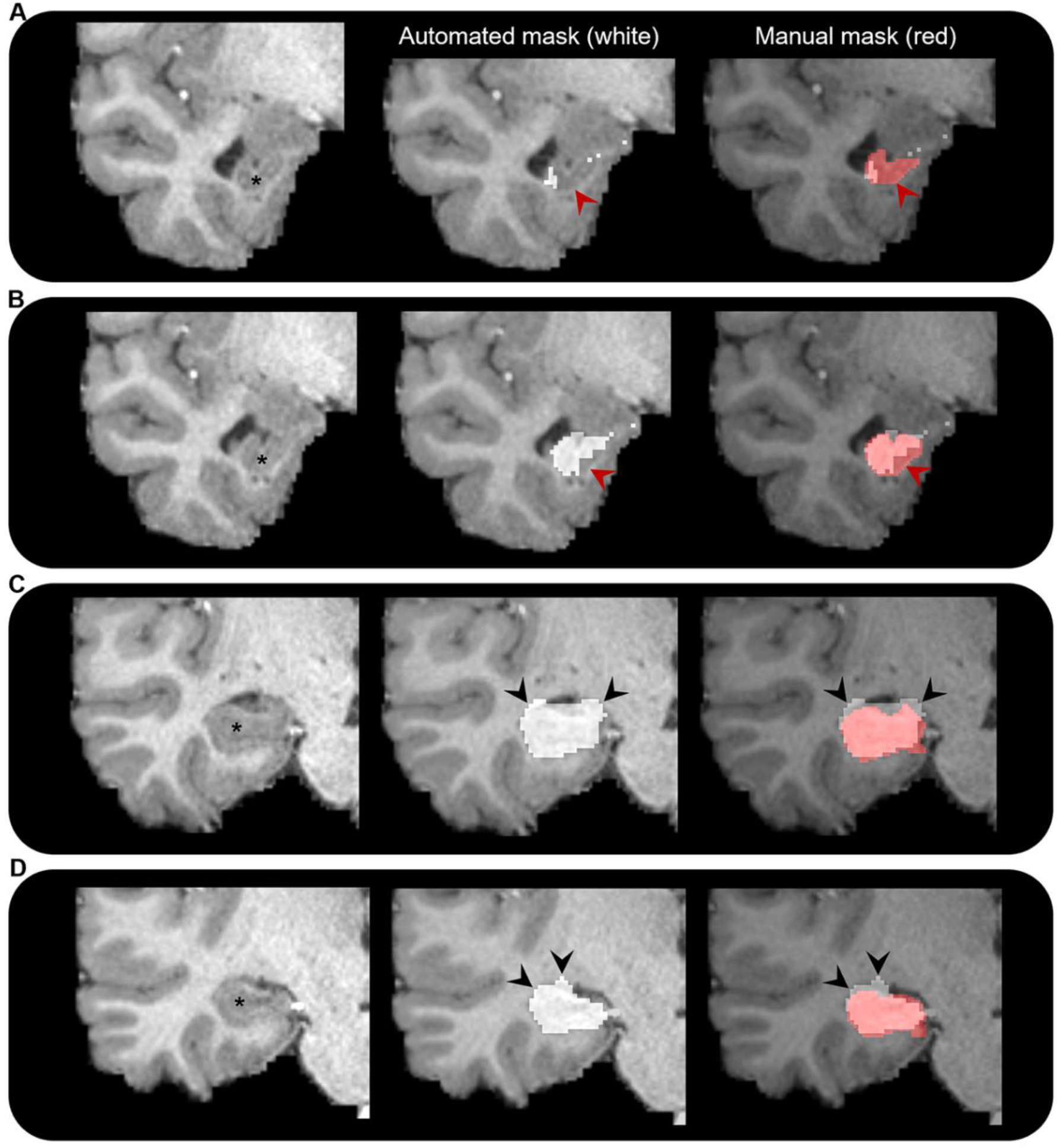
Comparison of automated and manual hippocampus segmentations. Representative examples of the automated hippocampus mask derived from the HCPMMP and the manually segmented hippocampus mask. We display examples from anterior (a) to posterior (d) portions of the hippocampal head. In each panel, we present a coronal slice of the T1-weighted image focussed on the right temporal lobe for a single participant (left; hippocampus indicated by *), the same image overlaid with the automated hippocampus mask derived from the HCPMMP (middle; white) and the same image overlaid with both the automated HCPMMP hippocampus mask (right; white) and the manually segmented hippocampus mask (transparent red). Note, in the anterior most slices (a and b) the automated mask does not cover the entire extent of the hippocampus (indicated by red arrows) and in more posterior slices (c and d) the automated mask often overextends across the lateral ventricle superior to the hippocampus and into the adjacent white matter (indicated by black arrows). Streamlines making contact with these erroneous portions of the automated hippocampus mask may lead to results that are biologically implausible.

**Fig. S7.**
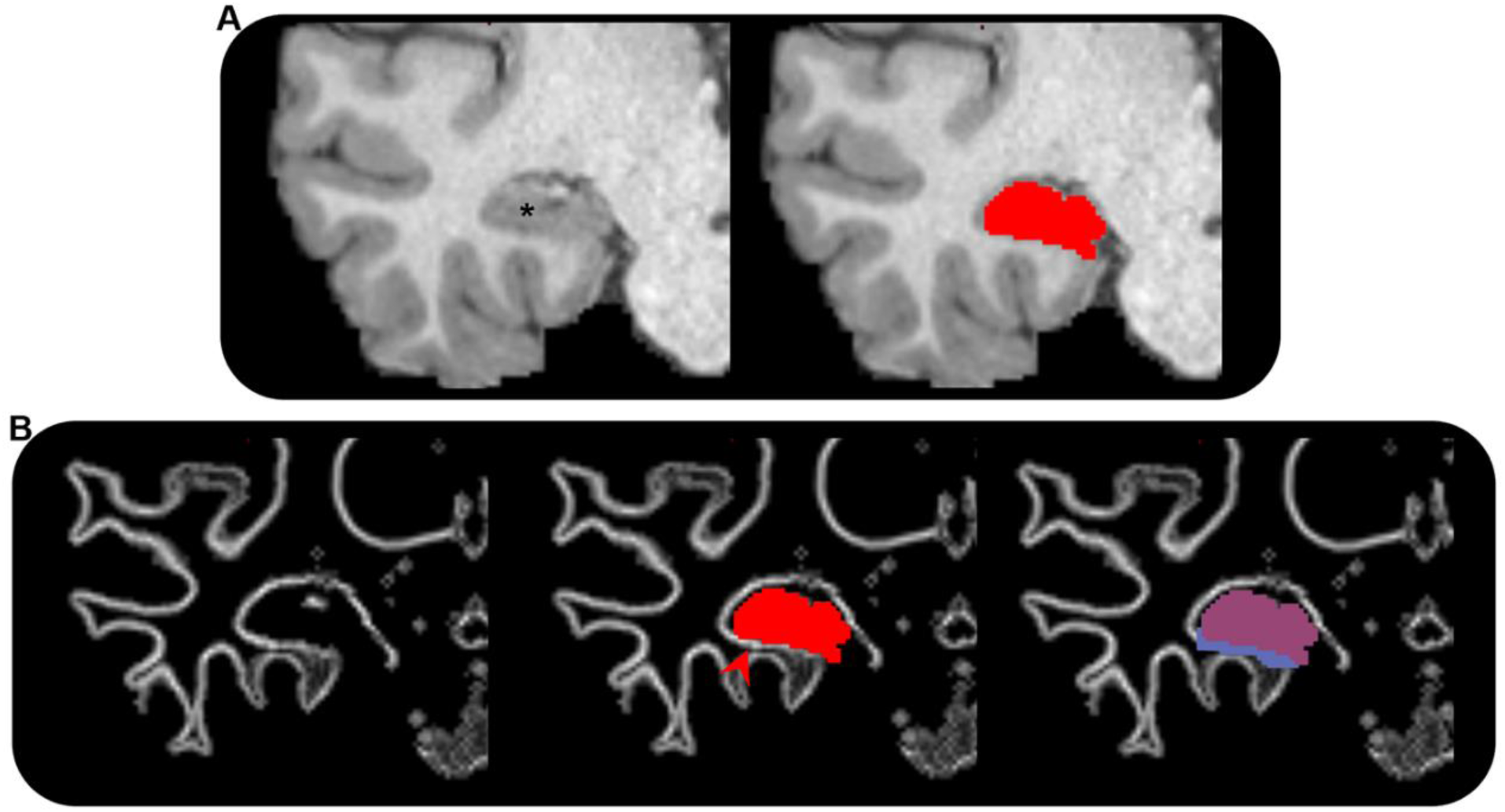
Adjustment of grey matter/white matter interface underlying the hippocampus. a. T1-weighted structural MRI scan showing the right medial temporal lobe (left; hippocampus indicated by *) in the coronal plane for a single participant and the manually segmented hippocampus mask (red) overlaid on the T1-weighted image (right). b. (left) The grey matter/white matter (gm-wm) interface (white line) showing the right medial temporal lobe in the coronal plane; (middle) the manually segmented hippocampus mask (red) overlaid on the gm-wm interface. Note: portions of detected gm-wm interface immediately inferior to the hippocampus lie outside of the hippocampus mask (indicated by red arrow). If left unchanged, the anatomically-informed tractography algorithm used in our study would terminate tracks as they reach this band, thus creating an erroneous band of track end-points in this region, introducing misleading results. (Right) The hippocampus mask (transparent red) and extended hippocampus mask (blue) overlaid on the gm-wm interface. Note: the extended hippocampus mask encompasses the gm-wm interface immediately inferior to the hippocampus. This allows streamlines to enter/leave the hippocampus here rather than terminate at the gm-wm interface (see Methods for detail).

**Fig. S8.**
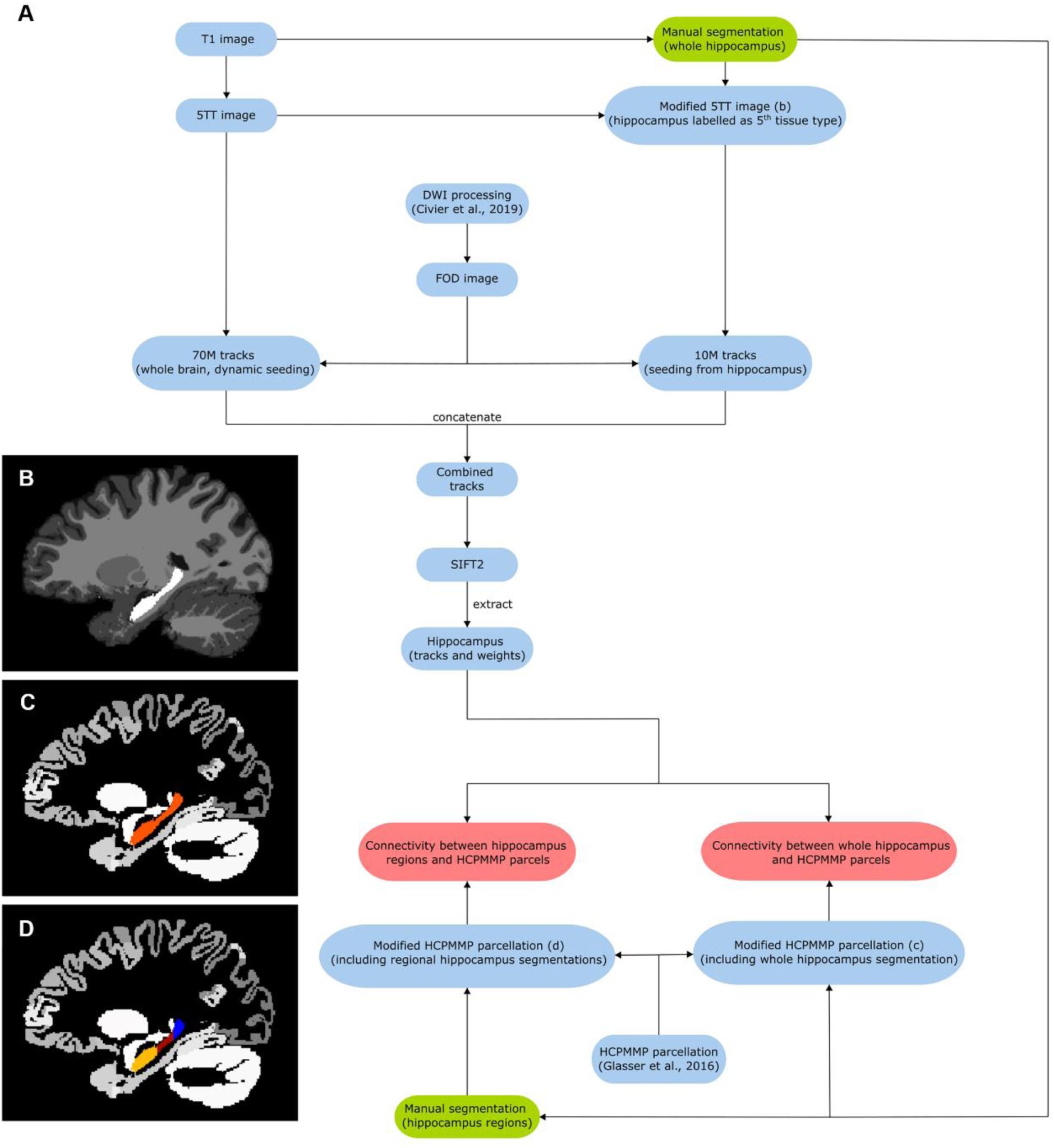
Analysis pipeline. a. Block diagram of the workflow. Green blocks indicate procedures that involved manual segmentation, red blocks indicate connectivity vectors/matrices from which connectivity measurements were obtained and blue blocks indicate intermediate images. b. Sagittal view of ‘modified 5TT’ image containing the manually segmented hippocampus labelled as ‘5th tissue type’ (i.e. no anatomical prior for tracking) shown in white. c. Sagittal view of modified parcellation image containing the whole-hippocampus mask (shown in orange). d. Sagittal view of modified parcellation image containing the regional hippocampus masks (head, body and tail shown in yellow, red and blue, respectively).

**Table S1.**
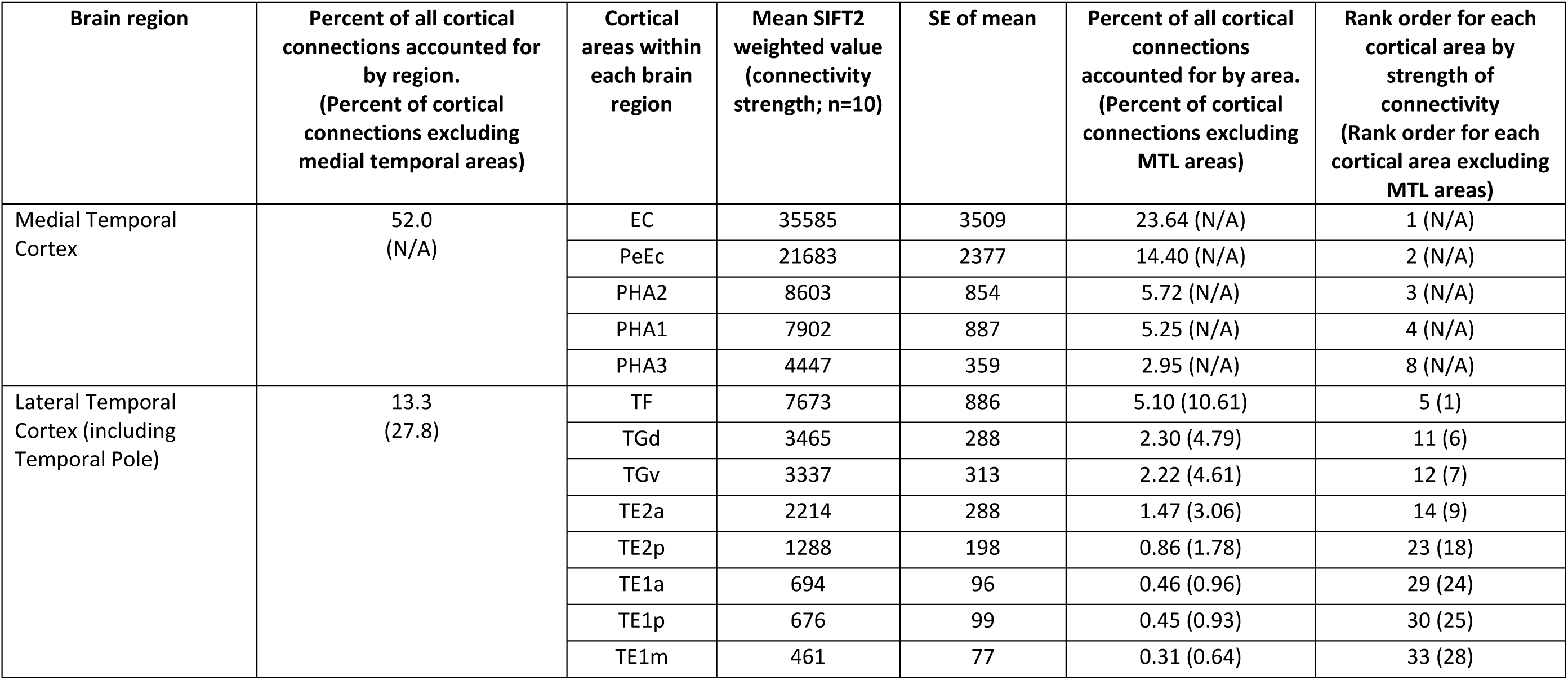
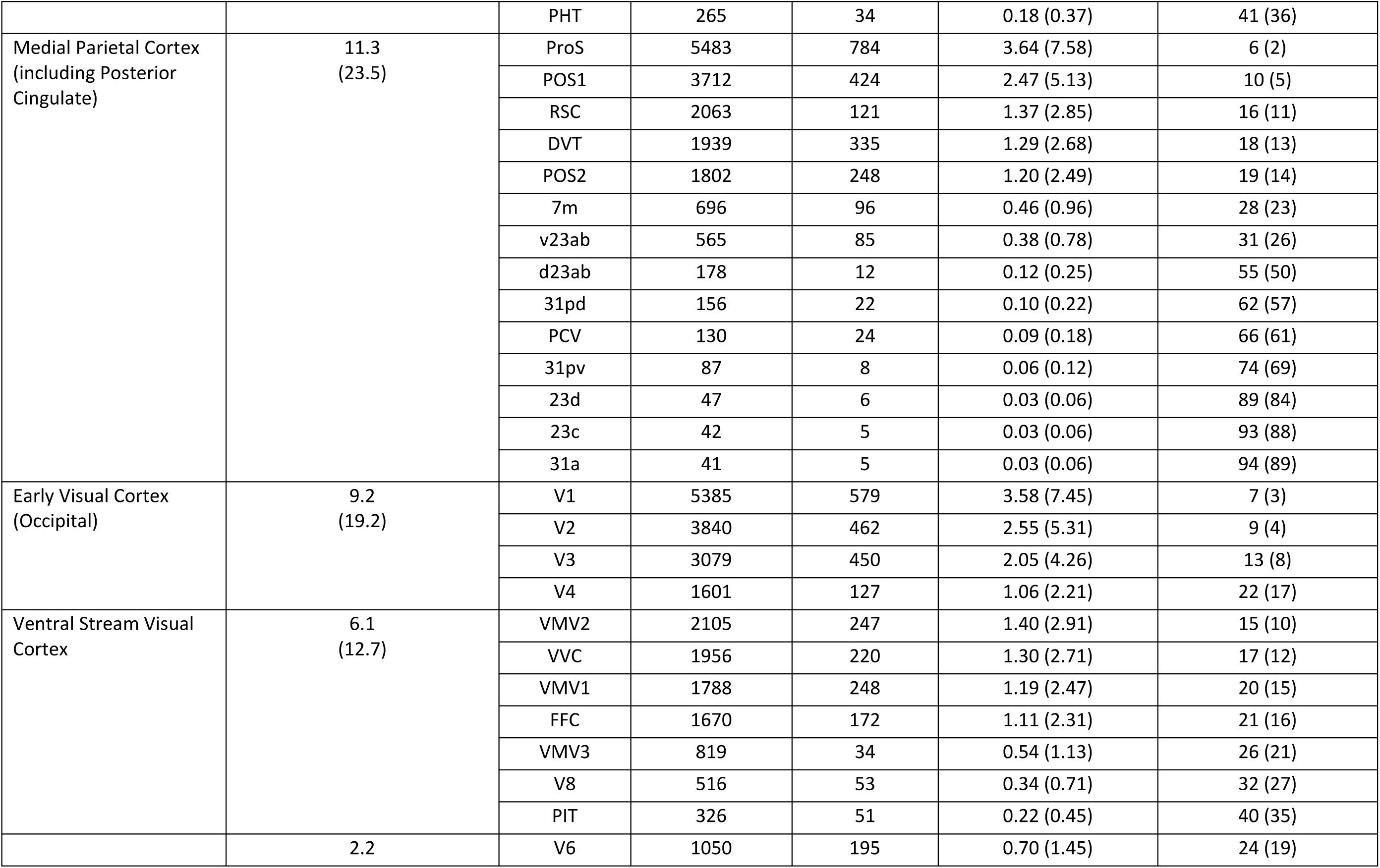
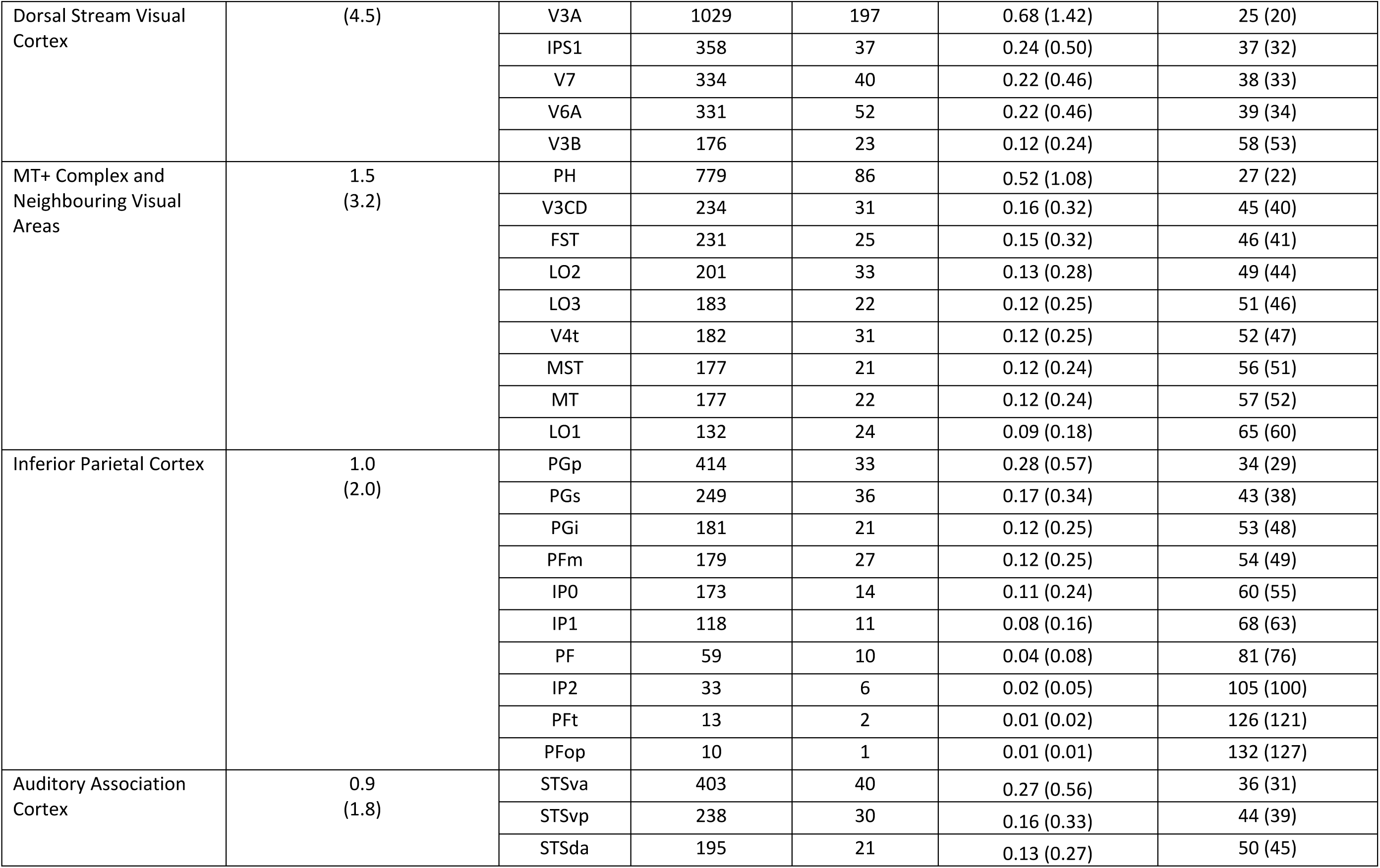
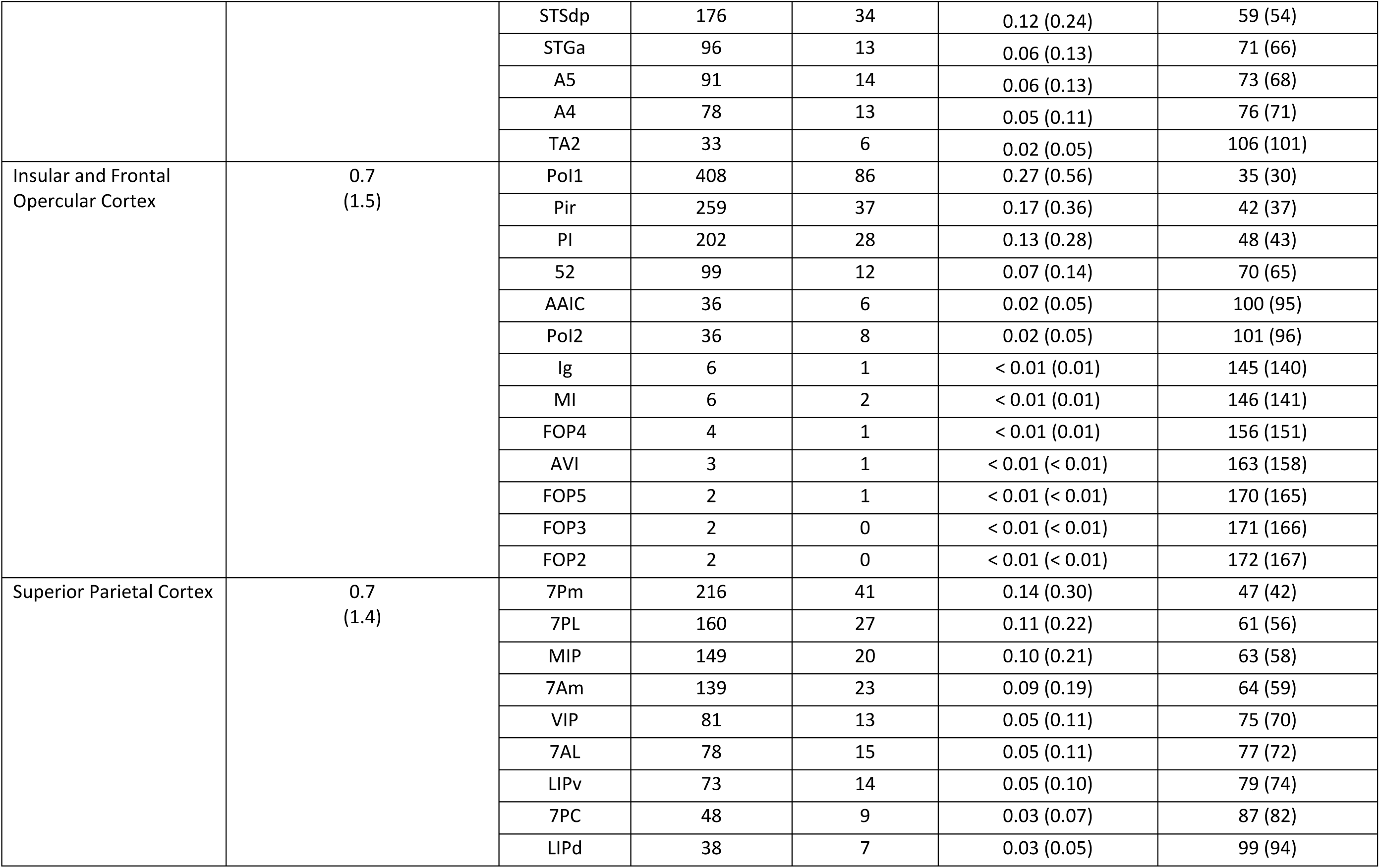
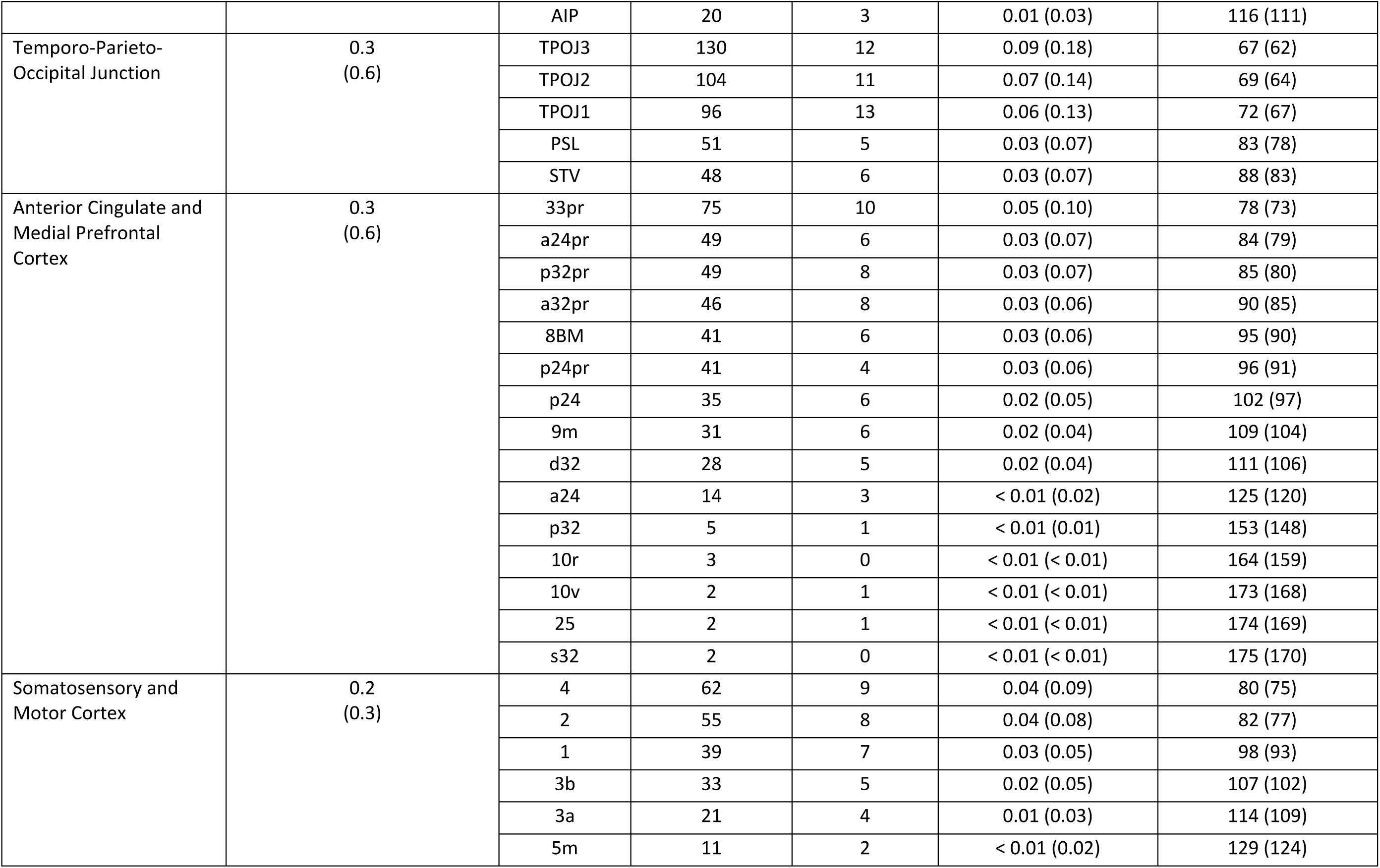
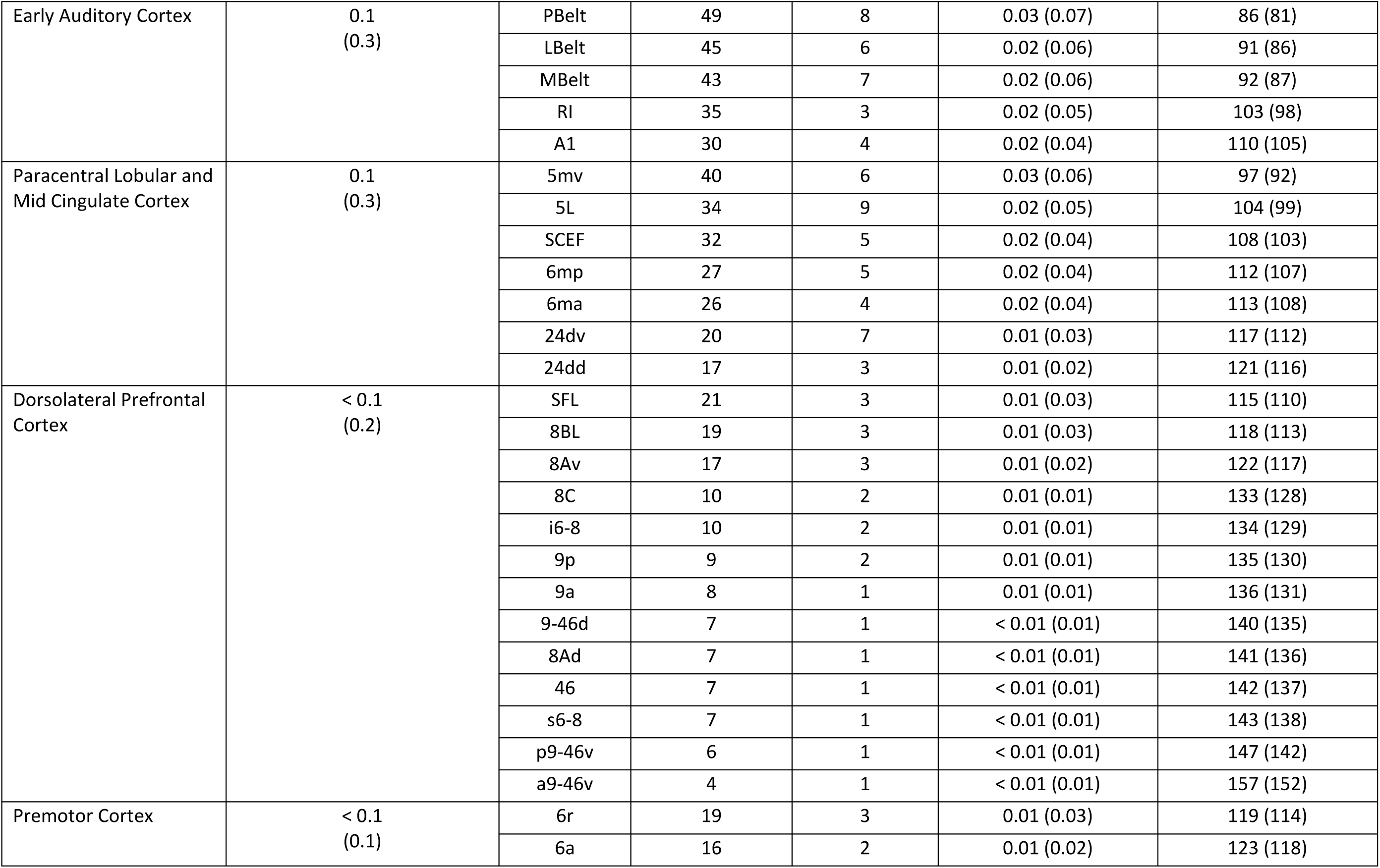
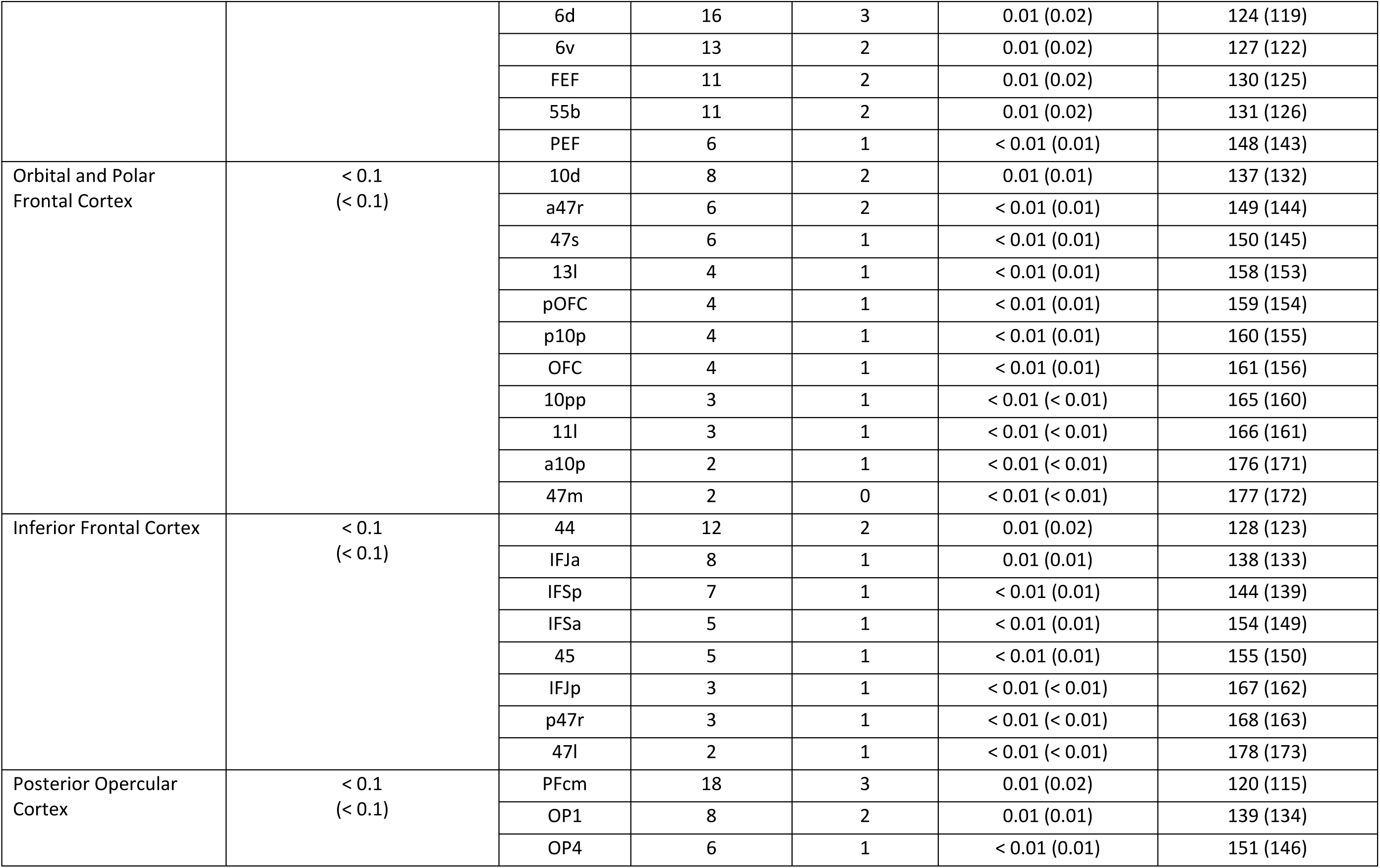
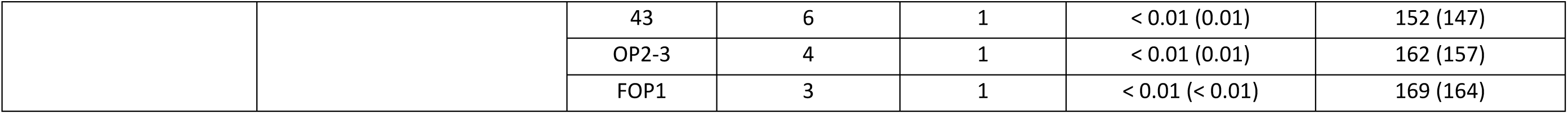
Connectivity values between the cortical mantle and the whole hippocampus. Connectivity values between the whole hippocampus and all cortical areas in the Human Connectome Project Multi-Modal Parcellation (HCPMMP) scheme are presented. Column 1 displays each broad brain region ordered by their strength of connectivity with the whole hippocampus. Column 2 displays the percent of all cortical connections accounted for by each region. Values in brackets indicate the percent of cortical connections accounted for by each region when excluding medial temporal lobe (MTL) areas. Column 3 presents the cortical areas located within each broad brain region ordered by their strength of connectivity with the whole hippocampus (abbreviations for all cortical areas are defined in Table S3). Column 4 displays the mean SIFT2 weighted value (connectivity strength) associated with each cortical area. Column 5 displays the standard error of the mean. Column 6 displays the percent of all cortical connections accounted for by each cortical area. Values in brackets indicate the percent of cortical connections accounted for by each cortical area when excluding MTL areas. Column 7 displays the rank order for each cortical area by strength of connectivity. Values in brackets indicate the rank order when excluding MTL areas.

**Table S2.** Connectivity between the cortical mantle and the head, body and tail of the hippocampus. Connectivity values between the head, body and tail of the hippocampus and all cortical brain areas included in the Human Connectome Project Multi-Modal Parcellation (HCPMMP) scheme are presented. Column 1 displays each cortical area ordered by strength of connectivity with the whole hippocampus (abbreviations for all cortical areas are defined in Table S3). Column 2 presents the broader brain region within which each cortical area is located. Column 3 displays the mean SIFT2 weighted value (connectivity strength) associated with each cortical area and the head of the hippocampus. Column 4 displays the associated standard error of the mean. Column 5 displays the mean SIFT2 weighted value (connectivity strength) associated with each cortical area and the body of the hippocampus. Column 6 displays the associated standard error of the mean. Column 7 displays the mean SIFT2 weighted value (connectivity strength) associated with each cortical area and the tail of the hippocampus. Column 8 displays the associated standard error of the mean.

| Cortical area | Location of area | Head |  | Body |  | Tail |  |
| --- | --- | --- | --- | --- | --- | --- | --- |
|  |  | Mean SIFT2 weighted value (n=10) | SE of Mean | Mean SIFT2 weighted value (n=10) | SE of Mean | Mean SIFT2 weighted value (n=10) | SE of Mean |
| EC | Medial Temporal Cortex | 16206 | 1927 | 12679 | 1357 | 6700 | 843 |
| PeEc | Medial Temporal Cortex | 11144 | 1414 | 7838 | 998 | 2700 | 366 |
| PHA2 | Medial Temporal Cortex | 2644 | 322 | 4545 | 566 | 1414 | 116 |
| PHA1 | Medial Temporal Cortex | 2287 | 299 | 3716 | 450 | 1899 | 243 |
| TF | Lateral Temporal Cortex | 3287 | 500 | 3452 | 410 | 934 | 144 |
| ProS | Medial Parietal Cortex (including Posterior Cingulate) | 716 | 116 | 1625 | 194 | 3143 | 553 |
| V1 | Early Visual Cortex (Occipital) | 567 | 70 | 1389 | 112 | 3429 | 489 |
| PHA3 | Medial Temporal Cortex | 1420 | 134 | 2277 | 236 | 750 | 76 |
| V2 | Early Visual Cortex (Occipital) | 428 | 62 | 1054 | 82 | 2358 | 376 |
| POS1 | Medial Parietal Cortex (including Posterior Cingulate) | 620 | 89 | 1299 | 112 | 1793 | 297 |
| TGd | Lateral Temporal Cortex | 1576 | 182 | 1446 | 117 | 443 | 47 |
| TGv | Lateral Temporal Cortex | 1589 | 207 | 1358 | 144 | 389 | 41 |
| V3 | Early Visual Cortex (Occipital) | 327 | 37 | 810 | 83 | 1942 | 357 |
| TE2a | Lateral Temporal Cortex | 754 | 132 | 1038 | 107 | 422 | 65 |
| VMV2 | Ventral Stream Visual Cortex | 354 | 68 | 860 | 111 | 891 | 97 |
| RSC | Medial Parietal Cortex (including Posterior Cingulate) | 401 | 55 | 733 | 52 | 928 | 107 |
| VVC | Ventral Stream Visual Cortex | 460 | 76 | 969 | 129 | 528 | 48 |
| DVT | Medial Parietal Cortex (including Posterior Cingulate) | 247 | 36 | 515 | 65 | 1178 | 253 |
| POS2 | Medial Parietal Cortex (including Posterior Cingulate) | 303 | 32 | 590 | 47 | 909 | 195 |
| VMV1 | Ventral Stream Visual Cortex | 245 | 47 | 644 | 115 | 899 | 110 |
| FFC | Ventral Stream Visual Cortex | 355 | 49 | 798 | 109 | 517 | 55 |
| V4 | Early Visual Cortex (Occipital) | 183 | 20 | 472 | 40 | 946 | 116 |
| TE2p | Lateral Temporal Cortex | 348 | 69 | 658 | 101 | 282 | 45 |
| V6 | Dorsal Stream Visual Cortex | 113 | 16 | 259 | 37 | 678 | 149 |
| V3A | Dorsal Stream Visual Cortex | 103 | 11 | 252 | 35 | 674 | 155 |
| VMV3 | Ventral Stream Visual Cortex | 127 | 16 | 306 | 28 | 386 | 23 |
| PH | MT+ Complex and Neighbouring Visual Areas | 136 | 18 | 338 | 44 | 305 | 40 |
| 7m | Medial Parietal Cortex (including Posterior Cingulate) | 120 | 17 | 251 | 26 | 325 | 69 |
| TE1a | Lateral Temporal Cortex | 209 | 42 | 329 | 39 | 156 | 23 |
| TE1p | Lateral Temporal Cortex | 146 | 27 | 334 | 46 | 196 | 32 |
| v23ab | Medial Parietal Cortex (including Posterior Cingulate) | 91 | 14 | 207 | 23 | 267 | 60 |
| V8 | Ventral Stream Visual Cortex | 72 | 14 | 178 | 20 | 265 | 33 |
| TE1m | Lateral Temporal Cortex | 121 | 32 | 232 | 35 | 108 | 14 |
| PGp | Inferior Parietal Cortex | 58 | 7 | 147 | 12 | 209 | 25 |
| PoI1 | Insular and Frontal Opercular Cortex | 110 | 21 | 174 | 36 | 124 | 31 |
| STSva | Auditory Association Cortex | 108 | 17 | 198 | 17 | 98 | 9 |
| IPS1 | Dorsal Stream Visual Cortex | 38 | 6 | 100 | 6 | 220 | 27 |
| V7 | Dorsal Stream Visual Cortex | 38 | 6 | 88 | 6 | 207 | 32 |
| V6A | Dorsal Stream Visual Cortex | 45 | 8 | 92 | 14 | 194 | 33 |
| PIT | Ventral Stream Visual Cortex | 41 | 8 | 116 | 26 | 168 | 25 |
| PHT | Lateral Temporal Cortex | 52 | 10 | 127 | 15 | 86 | 13 |
| Pir | Insular and Frontal Opercular Cortex | 148 | 27 | 81 | 10 | 29 | 4 |
| PGs | Inferior Parietal Cortex | 37 | 8 | 100 | 13 | 112 | 17 |
| STSvp | Auditory Association Cortex | 46 | 8 | 122 | 15 | 70 | 9 |
| V3CD | MT+ Complex and Neighbouring Visual Areas | 20 | 3 | 65 | 7 | 149 | 26 |
| FST | MT+ Complex and Neighbouring Visual Areas | 34 | 5 | 92 | 13 | 105 | 14 |
| 7Pm | Superior Parietal Cortex | 43 | 9 | 77 | 16 | 96 | 19 |
| PI | Insular and Frontal Opercular Cortex | 55 | 12 | 89 | 9 | 58 | 9 |
| LO2 | MT+ Complex and Neighbouring Visual Areas | 23 | 3 | 65 | 11 | 113 | 24 |
| STSda | Auditory Association Cortex | 54 | 10 | 90 | 9 | 51 | 5 |
| LO3 | MT+ Complex and Neighbouring Visual Areas | 21 | 3 | 66 | 9 | 97 | 16 |
| V4t | MT+ Complex and Neighbouring Visual Areas | 24 | 5 | 62 | 13 | 96 | 18 |
| PGi | Inferior Parietal Cortex | 27 | 5 | 85 | 14 | 70 | 8 |
| PFm | Inferior Parietal Cortex | 27 | 6 | 80 | 11 | 73 | 12 |
| d23ab | Medial Parietal Cortex (including Posterior Cingulate) | 28 | 4 | 62 | 5 | 88 | 8 |
| MST | MT+ Complex and Neighbouring Visual Areas | 21 | 3 | 71 | 10 | 86 | 14 |
| MT | MT+ Complex and Neighbouring Visual Areas | 21 | 3 | 64 | 9 | 92 | 15 |
| V3B | Dorsal Stream Visual Cortex | 13 | 2 | 43 | 4 | 120 | 21 |
| STSdp | Auditory Association Cortex | 37 | 12 | 84 | 13 | 55 | 10 |
| IP0 | Inferior Parietal Cortex | 18 | 2 | 56 | 3 | 99 | 12 |
| 7PL | Superior Parietal Cortex | 28 | 6 | 58 | 11 | 73 | 11 |
| 31pd | Medial Parietal Cortex (including Posterior Cingulate) | 25 | 5 | 55 | 6 | 76 | 14 |
| MIP | Superior Parietal Cortex | 17 | 4 | 52 | 8 | 80 | 10 |
| 7Am | Superior Parietal Cortex | 23 | 4 | 53 | 9 | 63 | 12 |
| LO1 | MT+ Complex and Neighbouring Visual Areas | 12 | 3 | 38 | 8 | 83 | 18 |
| PCV | Medial Parietal Cortex (including Posterior Cingulate) | 23 | 5 | 47 | 9 | 60 | 13 |
| TPOJ3 | Temporo-Parieto-Occipital Junction | 15 | 2 | 51 | 4 | 64 | 10 |
| IP1 | Inferior Parietal Cortex | 15 | 3 | 43 | 3 | 60 | 7 |
| TPOJ2 | Temporo-Parieto-Occipital Junction | 16 | 3 | 48 | 6 | 40 | 5 |
| 52 | Insular and Frontal Opercular Cortex | 18 | 2 | 41 | 3 | 40 | 8 |
| TPOJ1 | Temporo-Parieto-Occipital Junction | 17 | 3 | 46 | 5 | 34 | 6 |
| STGa | Auditory Association Cortex | 31 | 7 | 45 | 5 | 19 | 3 |
| A5 | Auditory Association Cortex | 19 | 3 | 39 | 4 | 32 | 7 |
| 31pv | Medial Parietal Cortex (including Posterior Cingulate) | 14 | 2 | 32 | 3 | 41 | 5 |
| VIP | Superior Parietal Cortex | 12 | 3 | 31 | 5 | 38 | 5 |
| 7AL | Superior Parietal Cortex | 10 | 2 | 30 | 5 | 38 | 8 |
| A4 | Auditory Association Cortex | 14 | 3 | 34 | 4 | 30 | 6 |
| 33pr | Anterior Cingulate and Medial Prefrontal Cortex | 14 | 2 | 28 | 4 | 33 | 5 |
| LIPv | Superior Parietal Cortex | 8 | 2 | 28 | 5 | 37 | 8 |
| 4 | Somatosensory and Motor Cortex | 8 | 2 | 28 | 5 | 26 | 5 |
| PF | Inferior Parietal Cortex | 11 | 3 | 27 | 5 | 21 | 3 |
| 2 | Somatosensory and Motor Cortex | 7 | 1 | 25 | 4 | 23 | 3 |
| PSL | Temporo-Parieto-Occipital Junction | 9 | 1 | 24 | 3 | 18 | 2 |
| a24pr | Anterior Cingulate and Medial Prefrontal Cortex | 10 | 2 | 18 | 2 | 21 | 3 |
| p32pr | Anterior Cingulate and Medial Prefrontal Cortex | 12 | 3 | 18 | 3 | 20 | 4 |
| PBelt | Early Auditory Cortex | 9 | 2 | 21 | 2 | 19 | 4 |
| 7PC | Superior Parietal Cortex | 6 | 2 | 20 | 4 | 22 | 4 |
| STV | Temporo-Parieto-Occipital Junction | 7 | 2 | 23 | 3 | 18 | 3 |
| 23d | Medial Parietal Cortex (including Posterior Cingulate) | 7 | 1 | 18 | 3 | 22 | 3 |
| a32pr | Anterior Cingulate and Medial Prefrontal Cortex | 11 | 3 | 18 | 3 | 17 | 3 |
| LBelt | Early Auditory Cortex | 7 | 1 | 19 | 2 | 19 | 3 |
| MBelt | Early Auditory Cortex | 6 | 1 | 19 | 2 | 17 | 4 |
| 23c | Medial Parietal Cortex (including Posterior Cingulate) | 6 | 1 | 15 | 2 | 21 | 3 |
| 8BM | Anterior Cingulate and Medial Prefrontal Cortex | 10 | 2 | 16 | 3 | 15 | 3 |
| p24pr | Anterior Cingulate and Medial Prefrontal Cortex | 7 | 1 | 15 | 2 | 19 | 3 |
| 31a | Medial Parietal Cortex (including Posterior Cingulate) | 7 | 2 | 14 | 1 | 20 | 3 |
| 5mv | Paracentral Lobular and Mid Cingulate Cortex | 5 | 1 | 15 | 2 | 20 | 4 |
| 1 | Somatosensory and Motor Cortex | 4 | 1 | 19 | 3 | 16 | 3 |
| LIPd | Superior Parietal Cortex | 4 | 1 | 14 | 3 | 20 | 4 |
| AAIC | Insular and Frontal Opercular Cortex | 12 | 4 | 17 | 3 | 7 | 1 |
| PoI2 | Insular and Frontal Opercular Cortex | 13 | 5 | 14 | 3 | 8 | 1 |
| p24 | Anterior Cingulate and Medial Prefrontal Cortex | 7 | 2 | 13 | 2 | 15 | 2 |
| RI | Early Auditory Cortex | 4 | 1 | 18 | 1 | 13 | 2 |
| 5L | Paracentral Lobular and Mid Cingulate Cortex | 5 | 1 | 12 | 3 | 17 | 4 |
| 3b | Somatosensory and Motor Cortex | 3 | 1 | 15 | 3 | 15 | 3 |
| TA2 | Auditory Association Cortex | 8 | 2 | 15 | 3 | 11 | 2 |
| IP2 | Inferior Parietal Cortex | 4 | 1 | 14 | 3 | 15 | 3 |
| SCEF | Paracentral Lobular and Mid Cingulate Cortex | 5 | 1 | 13 | 2 | 14 | 3 |
| 9m | Anterior Cingulate and Medial Prefrontal Cortex | 8 | 2 | 12 | 3 | 11 | 2 |
| A1 | Early Auditory Cortex | 4 | 1 | 12 | 1 | 13 | 3 |
| d32 | Anterior Cingulate and Medial Prefrontal Cortex | 7 | 1 | 10 | 2 | 11 | 2 |
| 6mp | Paracentral Lobular and Mid Cingulate Cortex | 3 | 1 | 12 | 3 | 12 | 2 |
| 6ma | Paracentral Lobular and Mid Cingulate Cortex | 3 | 1 | 10 | 2 | 13 | 2 |
| SFL | DorsoLateral Prefrontal Cortex | 4 | 1 | 7 | 1 | 10 | 2 |
| 3a | Somatosensory and Motor Cortex | 2 | 1 | 11 | 2 | 8 | 1 |
| AIP | Superior Parietal Cortex | 3 | 1 | 8 | 1 | 10 | 2 |
| 24dv | Paracentral Lobular and Mid Cingulate Cortex | 4 | 2 | 6 | 2 | 10 | 4 |
| 6r | Premotor Cortex | 3 | 1 | 9 | 1 | 6 | 1 |
| 8BL | DorsoLateral Prefrontal Cortex | 4 | 1 | 7 | 2 | 8 | 1 |
| PFcm | Posterior Opercular Cortex | 3 | 1 | 8 | 1 | 7 | 1 |
| 8Av | DorsoLateral Prefrontal Cortex | 3 | 1 | 7 | 1 | 7 | 1 |
| 24dd | Paracentral Lobular and Mid Cingulate Cortex | 2 | 0 | 6 | 1 | 8 | 2 |
| 6a | Premotor Cortex | 2 | 0 | 7 | 1 | 7 | 1 |
| 6d | Premotor Cortex | 2 | 1 | 8 | 2 | 6 | 1 |
| a24 | Anterior Cingulate and Medial Prefrontal Cortex | 3 | 1 | 5 | 1 | 6 | 2 |
| 6v | Premotor Cortex | 2 | 1 | 6 | 1 | 5 | 1 |
| PFt | Inferior Parietal Cortex | 2 | 1 | 7 | 1 | 4 | 1 |
| 44 | Inferior Frontal Cortex | 2 | 1 | 5 | 1 | 4 | 1 |
| FEF | Premotor Cortex | 2 | 1 | 5 | 1 | 5 | 1 |
| 5m | Somatosensory and Motor Cortex | 2 | 0 | 4 | 1 | 5 | 1 |
| 55b | Premotor Cortex | 2 | 0 | 5 | 1 | 4 | 1 |
| PFop | Inferior Parietal Cortex | 2 | 0 | 5 | 1 | 4 | 1 |
| 8C | DorsoLateral Prefrontal Cortex | 2 | 0 | 5 | 1 | 3 | 1 |
| i6-8 | DorsoLateral Prefrontal Cortex | 2 | 0 | 4 | 1 | 4 | 1 |
| 9p | DorsoLateral Prefrontal Cortex | 1 | 0 | 4 | 1 | 4 | 1 |
| 10d | Orbital and Polar Frontal Cortex | 2 | 1 | 3 | 1 | 3 | 1 |
| 9a | DorsoLateral Prefrontal Cortex | 1 | 0 | 3 | 1 | 4 | 1 |
| OP1 | Posterior Opercular Cortex | 1 | 0 | 4 | 1 | 2 | 1 |
| IFJa | Inferior Frontal Cortex | 2 | 0 | 3 | 1 | 3 | 1 |
| 9-46d | DorsoLateral Prefrontal Cortex | 1 | 0 | 2 | 0 | 4 | 0 |
| 8Ad | DorsoLateral Prefrontal Cortex | 1 | 0 | 2 | 0 | 4 | 1 |
| 46 | DorsoLateral Prefrontal Cortex | 1 | 0 | 3 | 1 | 3 | 1 |
| IFSp | Inferior Frontal Cortex | 1 | 0 | 3 | 1 | 2 | 0 |
| s6-8 | DorsoLateral Prefrontal Cortex | 1 | 0 | 2 | 0 | 4 | 1 |
| p9-46v | DorsoLateral Prefrontal Cortex | 1 | 0 | 2 | 1 | 3 | 1 |
| OP4 | Posterior Opercular Cortex | 1 | 1 | 3 | 1 | 2 | 0 |
| a47r | Orbital and Polar Frontal Cortex | 1 | 0 | 2 | 1 | 3 | 1 |
| 43 | Posterior Opercular Cortex | 1 | 0 | 3 | 0 | 2 | 0 |
| Ig | Insular and Frontal Opercular Cortex | 1 | 0 | 3 | 0 | 2 | 0 |
| MI | Insular and Frontal Opercular Cortex | 1 | 0 | 3 | 1 | 2 | 1 |
| 47s | Orbital and Polar Frontal Cortex | 1 | 0 | 3 | 1 | 2 | 1 |
| PEF | Premotor Cortex | 1 | 0 | 2 | 0 | 2 | 1 |
| p32 | Anterior Cingulate and Medial Prefrontal Cortex | 1 | 0 | 2 | 0 | 2 | 1 |
| IFSa | Inferior Frontal Cortex | 1 | 0 | 2 | 0 | 2 | 0 |
| 45 | Inferior Frontal Cortex | 1 | 0 | 2 | 0 | 2 | 0 |
| 13l | Orbital and Polar Frontal Cortex | 1 | 0 | 2 | 1 | 1 | 0 |
| pOFC | Orbital and Polar Frontal Cortex | 1 | 0 | 2 | 1 | 1 | 0 |
| FOP4 | Insular and Frontal Opercular Cortex | 1 | 0 | 2 | 0 | 2 | 0 |
| p10p | Orbital and Polar Frontal Cortex | 1 | 0 | 1 | 0 | 2 | 0 |
| OFC | Orbital and Polar Frontal Cortex | 1 | 0 | 2 | 1 | 1 | 0 |
| a9-46v | DorsoLateral Prefrontal Cortex | 1 | 0 | 2 | 0 | 1 | 0 |
| OP2-3 | Posterior Opercular Cortex | 1 | 0 | 2 | 0 | 1 | 0 |
| FOP1 | Posterior Opercular Cortex | 0 | 0 | 2 | 1 | 1 | 0 |
| IFJp | Inferior Frontal Cortex | 1 | 0 | 1 | 0 | 1 | 0 |
| p47r | Inferior Frontal Cortex | 1 | 0 | 1 | 0 | 1 | 0 |
| 10r | Anterior Cingulate and Medial Prefrontal Cortex | 1 | 0 | 1 | 0 | 1 | 0 |
| 10pp | Orbital and Polar Frontal Cortex | 1 | 0 | 1 | 0 | 1 | 0 |
| AVI | Insular and Frontal Opercular Cortex | 1 | 0 | 1 | 1 | 1 | 0 |
| 11l | Orbital and Polar Frontal Cortex | 1 | 0 | 1 | 0 | 1 | 0 |
| 10v | Anterior Cingulate and Medial Prefrontal Cortex | 1 | 0 | 1 | 0 | 1 | 0 |
| 47l | Inferior Frontal Cortex | 0 | 0 | 1 | 0 | 1 | 0 |
| a10p | Orbital and Polar Frontal Cortex | 0 | 0 | 1 | 0 | 1 | 0 |
| FOP5 | Insular and Frontal Opercular Cortex | 0 | 0 | 1 | 0 | 1 | 1 |
| 25 | Anterior Cingulate and Medial Prefrontal Cortex | 0 | 0 | 1 | 0 | 1 | 0 |
| FOP3 | Insular and Frontal Opercular Cortex | 0 | 0 | 1 | 0 | 1 | 0 |
| FOP2 | Insular and Frontal Opercular Cortex | 0 | 0 | 1 | 0 | 1 | 0 |
| 47m | Orbital and Polar Frontal Cortex | 0 | 0 | 1 | 0 | 1 | 0 |
| s32 | Anterior Cingulate and Medial Prefrontal Cortex | 0 | 0 | 1 | 0 | 0 | 0 |

**Table S3.**
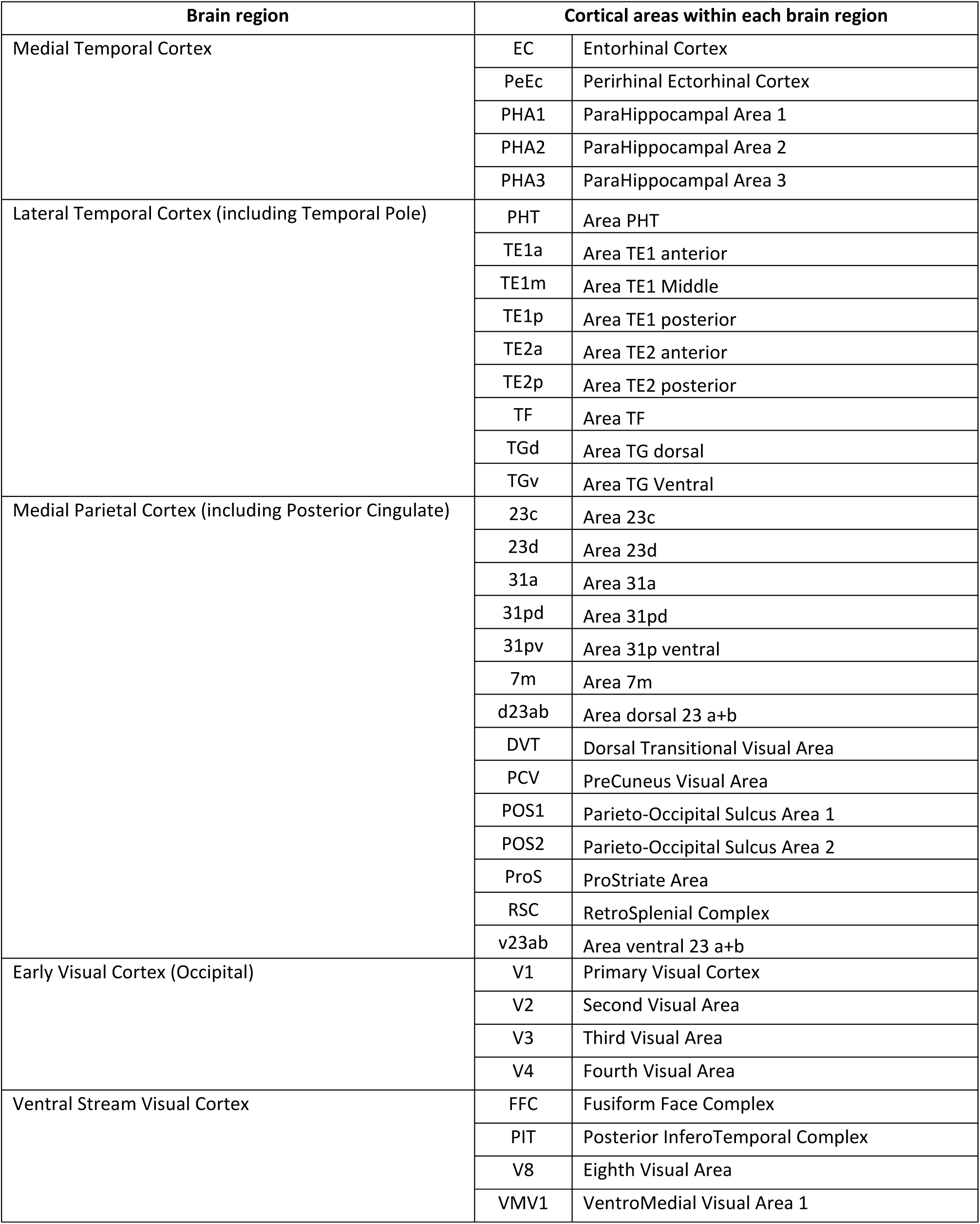
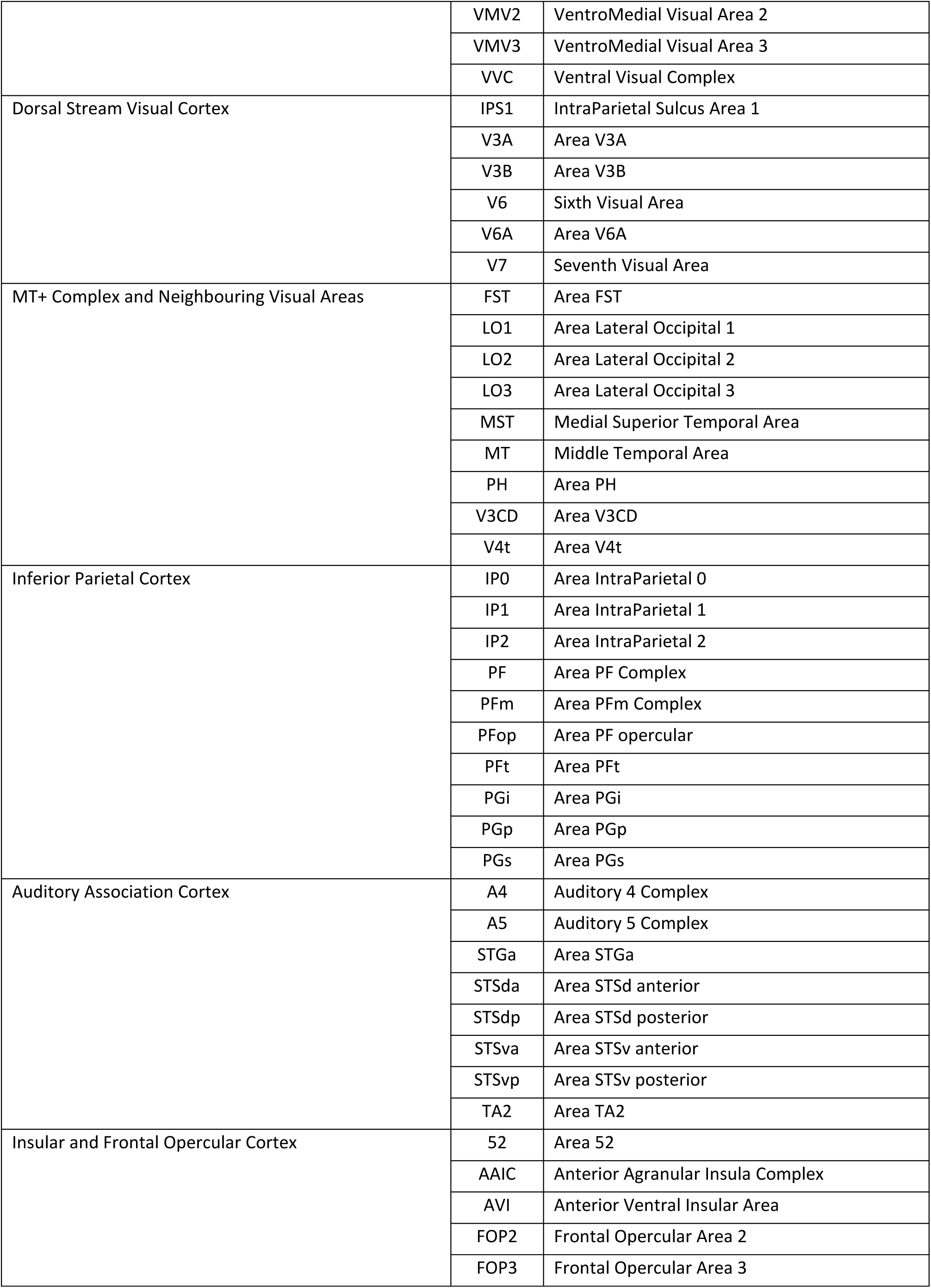
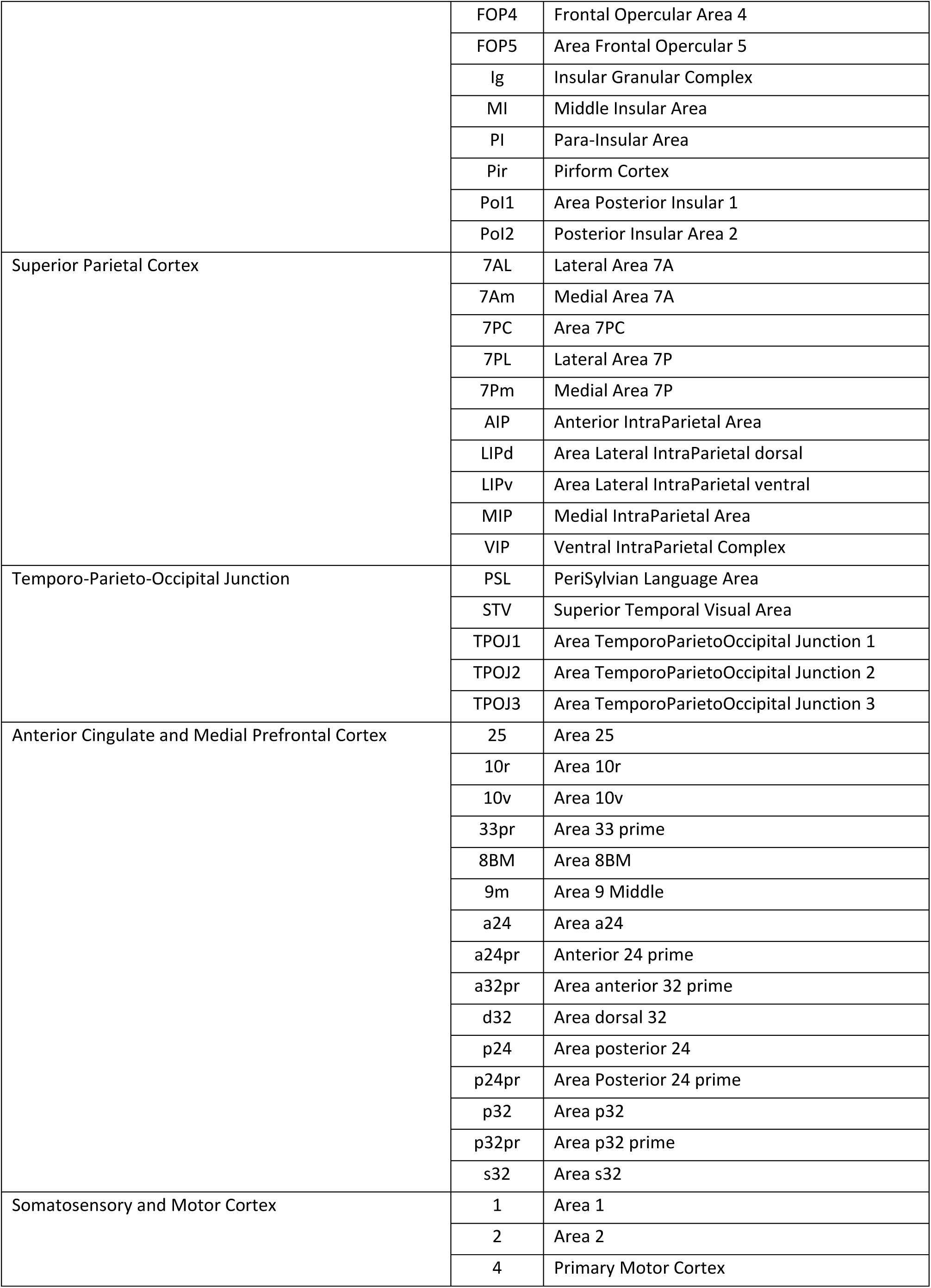
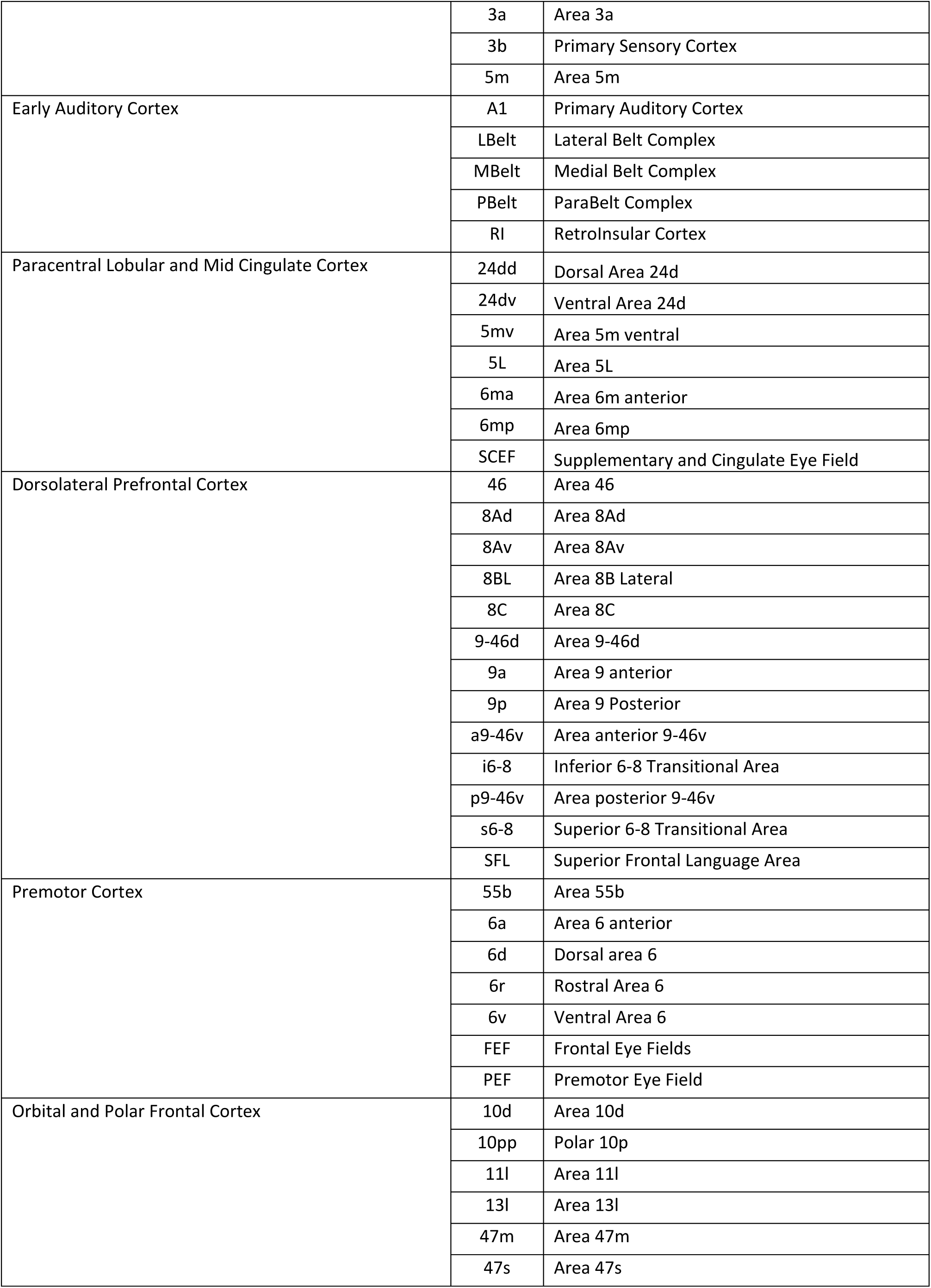
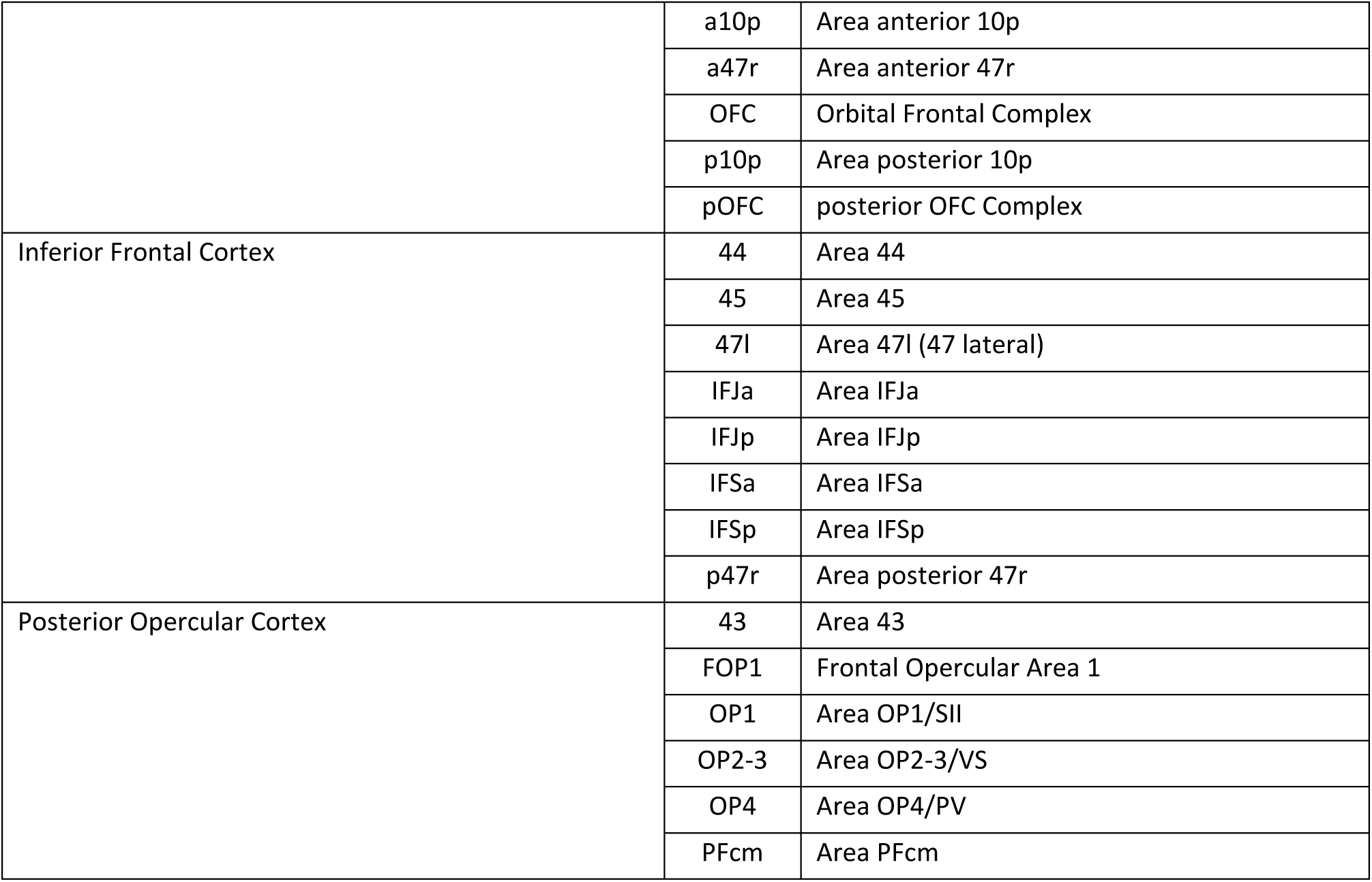
List of abbreviations for all cortical brain areas in the Human Connectome Project Multi-Modal Parcellation (HCPMMP) scheme.

## Footnotes

1 A preliminary version of this work was presented at the 30^th^ Annual Meeting of the International Society for Magnetic Resonance in Medicine (May 17, 2021).

## Notes

### Competing Interest Statement

The authors have declared no competing interest.

